# Reprogramming of alveolar macrophages by intestinal segmented filamentous bacteria protects mice from lethal bacterial pneumoniae following influenza infection

**DOI:** 10.1101/2025.01.23.634531

**Authors:** Vu L. Ngo, Carolin M. Lieber, Hirohito Abo, Michal Kuczma, Richard K. Plemper, Andrew T. Gewirtz

**Affiliations:** Center for Translational Antiviral Research, Georgia State University Institute for Biomedical Sciences, Atlanta GA 30303, USA

## Abstract

The most severe outcomes of respiratory viral infection (RVI) result from secondary bacterial infection, which RVI promotes via depletion of alveolar macrophages (AM). Colonization of the intestine by the common but non-ubiquitous commensal, segmented filamentous bacteria (SFB), reprograms AM to resist RVI-induced depletion. Hence, we examined if SFB against secondary infection by *S. pneumoniae*, *H. influenzae*, or *S. aureus,* following primary infection by influenza virus (IAV). Indeed, SFB colonization conferred strong post-IAV protection against these lethal bacterial pathogens. AM depletion and transplant studies indicated that SFB reprogramming these cells was necessary and sufficient for such protection. Assay of AM, *ex vivo*, from SFB-colonized mice argued their protection against secondary bacterial infection was not only due to their withstanding IAV-induced depletion. Rather, AM from SFB-colonized mice displayed complement-dependent increases in phagocytosis and killing of these bacteria. Furthermore, AM from SFB-colonized mice stably held their enhanced anti-bacterial phenotype even when transplanted into an inflamed interferon-rich post IAV-environment. Thus, SFB, and perhaps gut microbiota composition in general influences proneness to bacterial pneumonia, especially post-RVI.

**One Sentence summary:** SFB colonization stably changed the phenotype of alveolar macrophages resulting in sustained clearance of bacterial pathogens even amidst an inflamed interferon-rich immune suppressed lung.

## INTRODUCTION

A large portion of the mortality attributed to influenza virus (IAV) infection, and, to a less defined extent, respiratory viral infection (RVI) in general can be attributed to the secondary bacterial infections to which RVI greatly increases proneness (*1-3*). For example, histopathologic analysis of lung tissue specimens collected during the 1918 influenza pandemic found almost all lung specimens assayed from those deemed killed by the pandemic displayed visual evidence of robust infection by cocci, likely *S. pneumoniae*, or bacillus, likely *H. influenzae* (*4, 5*). These bacteria played a lesser role in influenza pandemics of the late 1950s in which many deaths were attributed to pneumonia driven by *S. aureus* (*6*). Secondary bacterial infections continue to be an intractable problem following infection by seasonal influenza viruses, which increase the risk of developing *S. pneumoniae*-induced pneumonia by 100-fold. Accordingly, secondary bacterial infections account for many RVI-related deaths, especially in elderly, despite frequent use of prophylactic antibiotics to stave them off.

Mechanisms by which RVI increase proneness to bacterial infection are complex but likely involves the absence of alveolar macrophages (AM), which are transiently depleted in an array of RVIs, including those caused by IAV, respiratory syncytial virus and SARS-CoV-2 (*7-9*). In support of this notion, pharmacologic depletion of AM, by itself, increases the proneness of mice to bacterial pneumonia, which, under such conditions, is not exacerbated by prior IAV infection (*10*). Mechanisms by which RVI results in AM depletion are poorly understood but plausibly results from caspase 3-mediated cell death triggered by high viral loads and pro-inflammatory cytokines (*11*).

We recently reported that colonization of the intestine by segmented filamentous bacteria (SFB), naturally acquired or exogenously administered, prevents RVI-induced AM depletion and, consequently, mitigates RVI severity (*11*). How SFB colonization prevents underlying RVI-induced depletion of AM is not fully defined but is a consequence of AM-intrinsic changes induced by this bacterium. More specifically, SFB colonization of the intestine results in AM becoming less prone to activation of pro-inflammatory gene expression while having greater capacity to disable IAV *ex vivo* due to enhanced complement production and phagocytosis. Such inflammatory anergy and enhanced antiviral function contribute to AM withstanding IAV-induced depletion *in vivo*. We reasoned that such changes in AM might also result in better protection against infection by respiratory bacterial pathogens (RBP), especially those which, most problematically, occur following RVI. We herein report observations that support this hypothesis. Specifically, we found that, even in the absence of RVI, SFB colonization conferred AM with greater ability to neutralize common RBP. Furthermore, SFB colonization made AM refractory to IAV-induced dysfunction. These SFB-induced impacts on AM resulted in stark protection against post-IAV bacterial infection.

## RESULTS

### SFB colonization protected mice from secondary or co-bacterial infection following primary IAV infection

Infection of the respiratory tract by an array of viruses, including IAV, results in depletion of alveolar macrophages (AM), thereby increasing proneness to bacterial pneumonia (*8, 12*). Colonization of the intestine by Segmented filamentous bacteria (SFB) prevents such depletion (*11*). Accordingly, we hypothesized that SFB might protect against post-IAV secondary bacterial infection. To test this hypothesis, we first established experimental conditions in mice not colonized with SFB (i.e. SFB^-^ mice), in which IAV infection depleted AM but was, by itself, non-lethal. C57BL/6 mice were intranasally administered 2009 pandemic A/CA/07/2009 (H1N1) IAV, herein referred to as CA09, which readily replicates in mice without prior species adaptation (*13*). A relatively moderate inoculum, namely 500 TCID_50_ units, resulted in a more than 80% reduction in AM levels but was uniformly lethal, thus precluding administration of a secondary bacterial challenge **(Fig S1)**. Reducing the inoculum by 10- and 100-fold resulted in more moderate levels of AM depletion at day 4 post infection, 54% and 40% respectively, and caused disease symptoms, including weight loss, but not death. Thus, we chose the former, 50 TCID_50_, for further study.

SFB**^-^** mice were orally administered vehicle or SFB and, thereafter, referred to as SFB^-^ or SFB**^+^** mice. Seven days later, SFB**^-^** and SFB**^+^** mice were inoculated with CA09 or mock-infected, followed by nasal administration of *S. pneumoniae* (10^7^ CFU) 4 days post infection (dpi). In mock-infected mice, *S. pneumoniae* was uniformly non-lethal irrespective of SFB although, compared to SFB**^-^** mice, SFB**^+^** mice displayed less weight loss and exhibited a more than one order of magnitude reduction in lung CFU **(Fig 1A)**, suggesting that SFB gut colonization conferred some protection against this respiratory pathogen. An analogous, albeit much more striking, pattern was observed when *S. pneumoniae* was administered following IAV infection. Specifically, CA09/*S. pneumoniae* infection resulted in 100% and 0% lethality in SFB**^-^** and SFB**^+^** mice, respectively. Such lethality, which occurred 5-6 days post *S. pneumoniae* administration, was, on day 4, associated with stark gross and histopathologically evident lung injury that was not observed in CA09/*S. pneumoniae*-infected SFB**^+^** mice. The potentiation of *S. pneumoniae*-induced disease by prior CA09 infection was associated with a one order of magnitude increases in lung *S. pneumoniae* CFU on day 4. In contrast, CA09 infection did not increase *S. pneumoniae* CFU in SFB**^+^** mice, resulting in starkly lower levels of *S. pneumoniae* CFU levels in lung, BALF, blood, and spleen **(Fig S2A)**. Similar results were obtained in secondary bacterial infection models in which CA09 inoculation was followed by *Hemophilus influenzae* (*H. influenzae*) and Methicillin-resistant *Staphylococcus aureus* (*S. aureus*) **(Fig 1B)**. These data support our hypothesis that SFB colonization protects against secondary bacterial infection.

**Figure 1.**
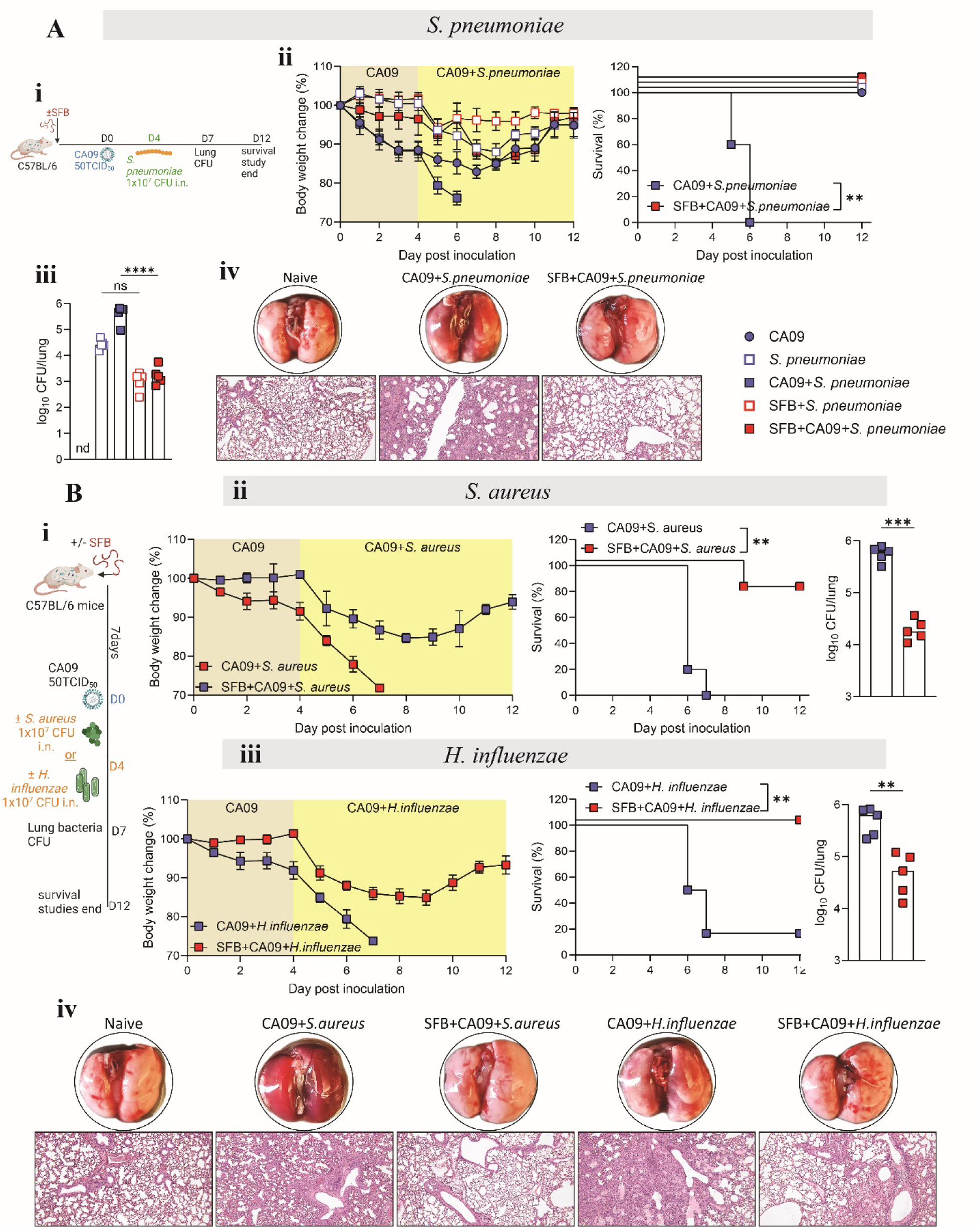
SFB protected mice from secondary/co-bacterial infection after primary influenza virus infection. **(A) (i)** Experimental scheme: Three-week-old C57BL/6 SFB^-^mice were orally administered with SFB. Seven day later, mice were inoculated with 50 TCID_50_ of CA09. On day four post-CA09 inoculation, mice were intranasally administered vehicle (PBS) or 1x10^7^ PFU of *S. pneumoniae.* **(ii)** Body weight and survival rate were monitored daily. **(iii)** Day 4 quantiation of lung bacterial CFU, and **(iv)** H&E stained lung sections. **(B) (i)** Experimental scheme: three-week-old C57BL/6 SFB^-^mice were colonized or not with SFB, and inoculated with CA09 as in (A). Four-day post-virus inoculation, mice were intranasally administered vehicle (PBS) or 1x10^7^ PFU of *S. aureus*, or *H. influenzae*. **(ii & iii)** Body weight, survival, and lung CFU of mice inoculated with *S. aureus* **(ii),** or *H. influenzae* **(iii)**. **(iv)** H&E stained lung sections. All experiments used five animal per group (n=5). Data are representative of two independent experiments, yielding an identical pattern of results. Results are shown as mean SD. Statistical analysis: lung bacterial burden: One-way ANOVA or Student’s t test. Survival: log-rank Mental-Cox Test. *p < 0.05, ** p < 0.01, **** p < 0.0001, ns not significant.

We hypothesized that the reduced bacterial loads in SFB^+^ mice reflected increased clearance of the bacterial pathogen. To probe this notion, we re-analyzed previously generated RNA-seq data from whole lungs of mice administered SFB and/or CA09, assayed at the time that approximately corresponds to the secondary bacterial infection administered herein (*13*). We found that SFB gut colonization caused a strong post-CA09 increase in lung expression of genes involved in mucosal clearance of bacterial pathogens **(Fig S2B)**. To directly test this notion, we performed post-CA09 *S. pneumoniae* infection using only 500 CFU and measured bacteria that remained two hours later. Mock-infected and SFB**^+^** CA09-infected mice fully cleared this bacterial inoculum. In contrast, 80% of the live inoculum was recovered in CA09-infected SFB**^-^** mice. These results indicate that SFB colonization enhanced bacterial clearance after CA09 infection.

### SFB-mediated protection against secondary bacterial infections in mice required alveolar macrophages

We hypothesized that AM mediated the resistance of SFB**^+^** mice to secondary bacterial infection. In accord with this notion, *S.pneumoniae* infection, by itself, modestly reduced AM levels and potentiated the much greater loss of these cells that occurred following CA09 infection **(Fig 2A)**. In stark contrast, AM levels of SFB**^+^** mice were similar to that of naïve mice irrespective of CA09 and *S. pneumoniae.* The temporal window of protection against secondary infection conferred by SFB also aligned it being mediated by AM. Specifically, SFB strongly protected against AM depletion and disease when *S. pneumoniae* was administered 2 or 10 dpi with CA09 **(Fig S3A)** but differences between SFB**^-^** and SFB**^+^** mice were modest 30 and 90 dpi, at which time the depleted AM would have been self-renewed or replaced by recruited monocytic precursors **(Fig S3B)** (*14*). Depletion of AM also supported their involvement in mediating SFB’s protection against *S. pneumoniae.* Specifically, administration of clodronate liposomes, which specifically deplete AM without altering levels of other lung immune cells (*11*), recapitulated the increased proneness to *S. pneumoniae* infection exhibited by CA09-infected mice as assessed by CFU, weight loss, and survival **(Fig 2B)**. Moreover, clodronate liposomes rendered SFB**^-^** and SFB**^+^** mice equally prone to *S. pneumoniae* infection. Furthermore, the relative resistance of SFB**^+^** mice to CA09/*S. pneumoniae* infection was eliminated by clodronate liposome treatment, assessed by CFU, weight loss, and mortality **(Fig 2C)**. These results suggest that the resistance of AM depletion following CA09 infection exhibited by SFB**^+^** mice was germane to their resistance to *S. pneumoniae* infection.

**Figure 2.**
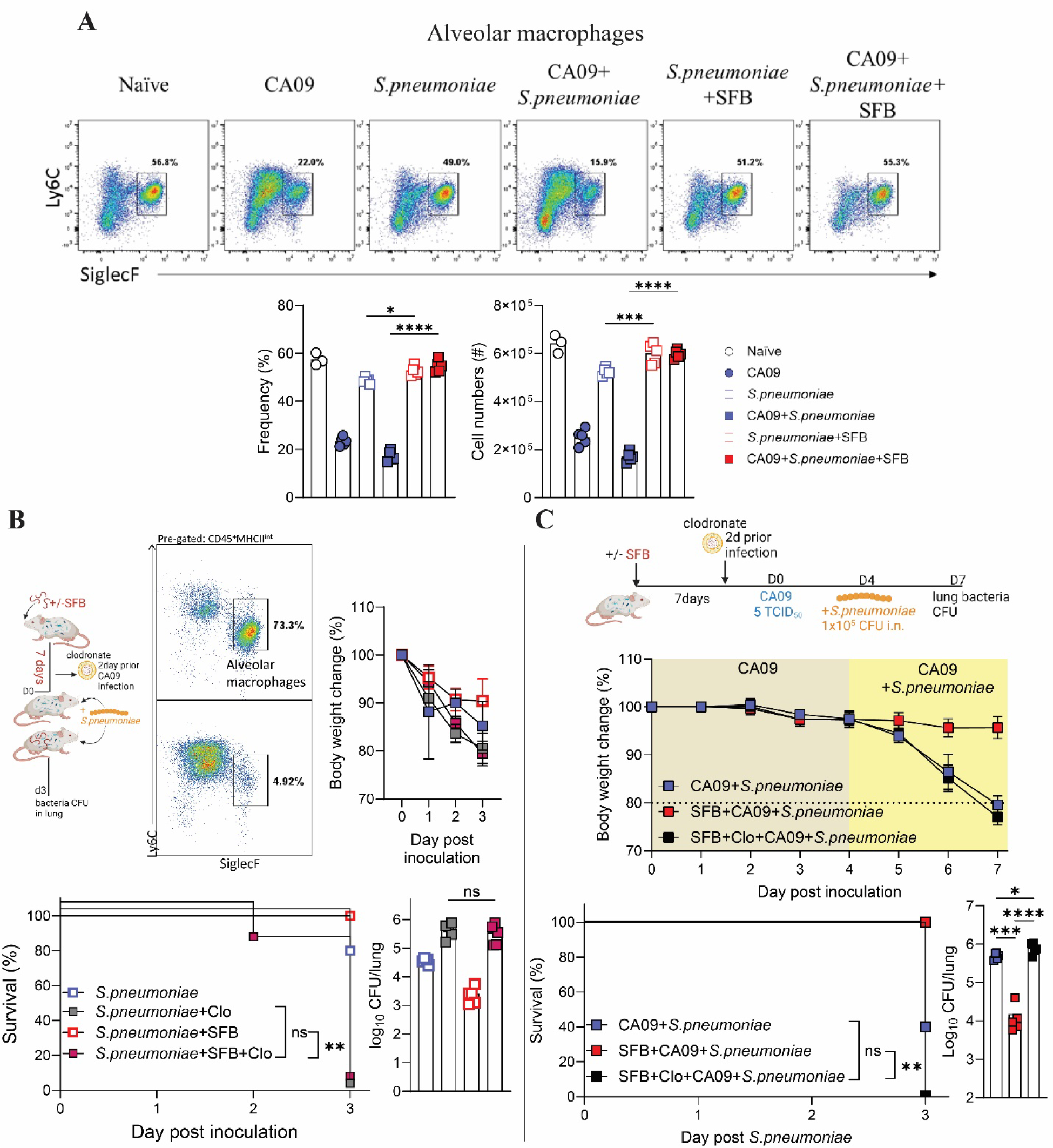
Resistance of SFB-colonized mice to secondary/co-bacterial infection required alveolar macrophages. **(A)** Three-week-old C57BL/6 SFB^-^mice, colonized with SFB or not for seven days prior and were inoculated with CA09. Four-days post virus inoculation, mice were inoculated intranasally with 1x10^7^ PFU of *S. pneumoniae*. Mice were euthanized four days later, and lung assayed by flow cytometry. Representative FACs plot, frequencies, and cell numbers of alveolar macrophages. **(B)** SFB^-^mice, colonized with SFB seven days prior or not, were nasally administered empty liposomes or clodronate liposomes two day prior to *S. pneumonaie* inoculation. Mice were than euthanized on day four post bacterial inoculation. FACs graphical representation lung AM after clodronate treatment, body weight, survival rate and lung bacterial burden were measured. **(C)** SFB^-^mice, colonized with SFB seven days prior or not, were nasally administered empty liposomes or clodronate liposomes two day prior to CA09 inoculation. Four-day post-virus inoculation, mice were inoculated with 1x10^7^ PFU of *S. pneumoniae* intranasally. Mice were euthanized on day four post bacterial inoculation. Body weight, survival rate, and lung bacterial burden were measured. All experiments used five animal per group (n=5). Data are representative of two independent experiments, yielding an identical pattern of results. Results are shown as mean SD. Statistical analysis: body weight: two-way ANOVA, lung bacterial burden: One-way ANOVA or Student’s t test. Survival: log-rank Mental-Cox Test. *p < 0.05, ** p < 0.01, *** p < 0.001, **** p < 0.0001, ns not significant.

### SFB reprogramming of alveolar macrophages enhances phagocytosis-mediated killing of bacteria

We presume that AM being present in greater numbers in SFB^+^ mice post-CA09 infection likely contributed to protection against *S. pneumoniae* infection. Yet, we envisaged that additional mechanisms may also be involved. More specifically, we hypothesized that AM derived from SFB**^+^** mice (hence forth referred to as SFB**^+^** AM) are intrinsically more adept at neutralizing IAV than SFB**^-^** AM led us to hypothesize that SFB**^+^** AM not only better cope with the CA09 challenge but may also have greater capacity per AM to eliminate bacteria they encounter. Re-analysis of our previously reported RNA seq data supported this notion.

Specifically, untargeted comparison of gene expression of SFB**^-^** and SFB**^+^** AM by Gene Set Enrichment Analysis found that genes involved in phagocytosis, complement activation, and secretion of antibacterial mediators were upregulated in SFB**^+^** AM **(Fig S4A)**. To probe the functional consequences of this observation, we isolated AM from SFB**^-^** and SFB**^+^** mice 4 dpi with CA09 via cell sorting, enabling us to expose equal numbers of SFB**^-^** and SFB**^+^** AM to relatively modest bacterial challenges, namely *S. pneumoniae*, *S. aureus*, or *H. influenzae* at a multiplicity of infection (MOI) of 20 per cell. Quantitation of CFU that remained in AM supernatants 60 minutes later indicated that SFB**^+^** AM were more capable to remove bacteria from their surroundings than SFB**^-^** AM **(Fig. 3ii)**. This pattern was more pronounced in AM isolated from CA09-infected mice (supernatants of SFB**^-^**CA09**^+^** AM contained 15-fold higher CFU than those of SFB**^+^**CA09**^+^** AM). In contrast, quantitation of intracellular CFU in AM at this time revealed more bacteria inside SFB**^+^** AM **(Fig S4D)**. Such rapid removal of bacteria from the suspension by SFB**^+^** AM, combined with increased intracellular CFU AM, suggested greater phagocytotic activity by SFB**^+^** AM. In accord with this notion SFB**^+^** AM exhibited greater ability to internalize fluorescent particles that had been coated with *S. aureus* antigens **(Fig S4B)**.

**Figure 3.**
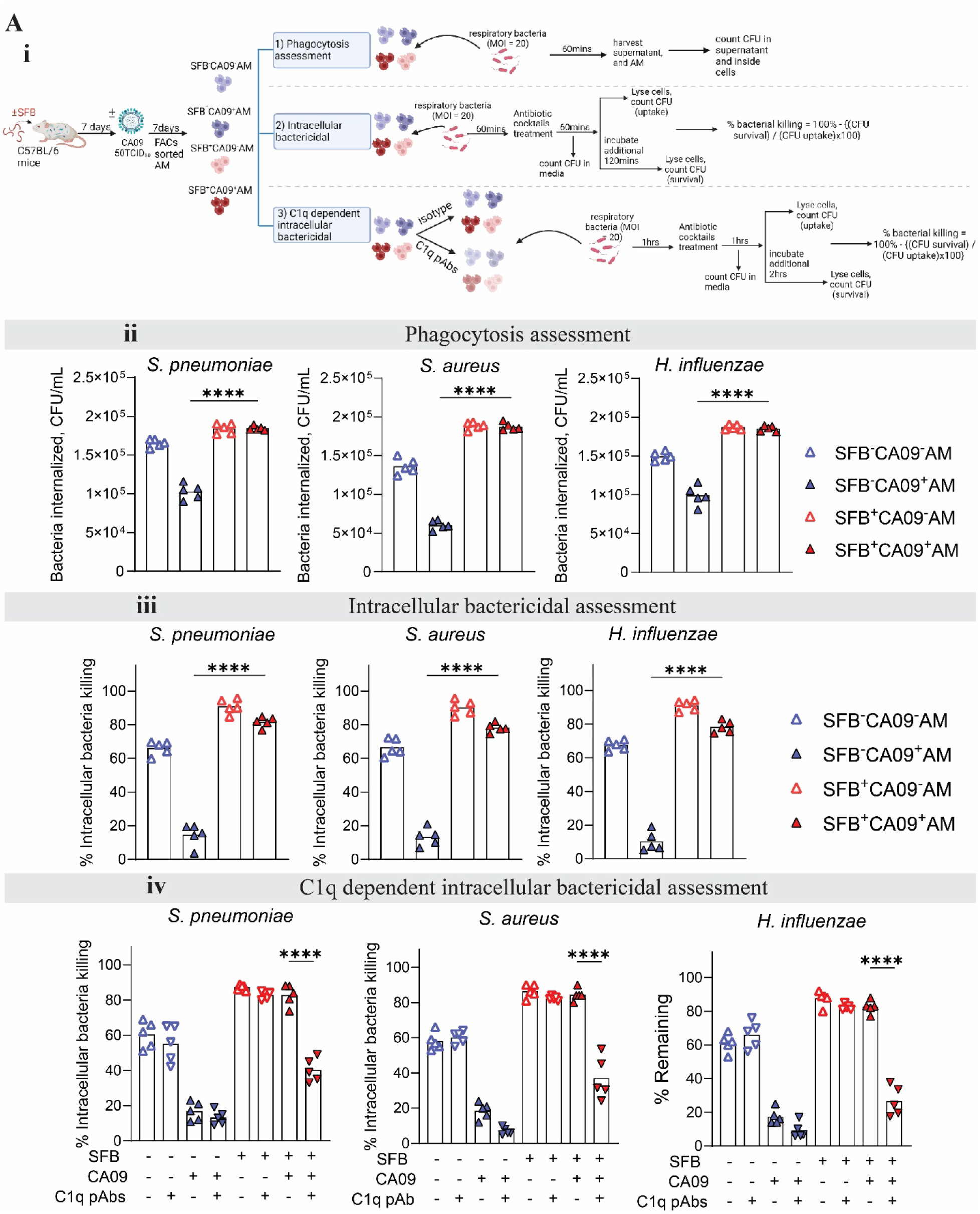
SFB^+^AM exhibited enhanced phagocytosis and C1q-depedent bactericidal activities. **(i)** Schematic design for *ex vivo* studies of AM. SFB^-^mice, colonized with SFB or not seven days prior, were inoculated with CA09. Five days post CA09 inoculation, SFB^-^CA09^-^AM, SFB^-^ CA09^+^AM, SFB^+^CA09^-^AM, and SFB^+^CA09^+^AM were FACs-sorted for downstream experiments. **(ii)** SFB moderately enhanced AM phagocytosis of bacteria *ex vivo*. FACs sorted AM were incubated with either *S. pneumoniae*, or *S. aureus*, or *H. influenzae* for sixty minutes. After incubation, supernatant was harvested, and CFU of bacterial remaining in supernatant were assay. Bacterial internalized were calculated as #original bacterial incubated with AM – bacterial remaining in the supernatant. **(iii)** SFB increases AM bactericidal *ex vivo*. FACs sorted AM were incubated with different bacteria and subjected to antibiotic protection assay and results were display as % intracellular bacterial killing. **(iv)** SFB^+^AM enhances phagocytosis and bactericidal require C1q complements. FACs-sorted AM were incubated with or without C1q-a/b/c neutralization or isotype antibodies for one hour. After incubation, supernatants were removed and replaced with new media. AM were then incubated with indicated bacteria and subjected to antibiotic protection as in (iii), and results are shown as % intracellular bacterial killing. n = 5 per group. Data are representative of two independent experiments, yielding an identical pattern of results. Results are shown as mean SD. Statistical analysis: Student’s t test (ii). One-way ANOVA (iii, iv). *p < 0.05, ** p < 0.01, *** p < 0.001, **** p < 0.0001, ns not significant.

SFB**^+^** AM also exhibited increased ability to kill the internalized bacteria. This difference was modest in absence of IAV infection. Specifically, relative to SFB**^-^** AM, SFB**^+^** AM began the assay with 2-fold higher levels of live intracellular bacteria (from greater phagocytosis) but nonetheless, over the next 2 hours, neutralized a greater proportion of them **(Fig 3iii, and Fig S4D)**. Such differential inactivation was more pronounced in AM isolated from CA09-infected mice. Specifically, SFB**^-^**CA09**^+^** AM neutralized only a small percentahe of the few bacteria they had internalized. In contrast, SFB**^+^**CA09**^+^** AM neutralized almost as well as SFB**^+^**CA09**^-^** AM and better than AM from naïve mice (i.e. SFB**^-^**CA09**^-^** AM). Thus, SFB colonization enhanced the ability of AM to neutralize bacterial pathogens and, furthermore, made AM largely refractory to IAV-induced AM dysfunction. We hypothesized a role for complement in mediating these differences between SFB**^-^** and SFB**^+^** AM. Measure of complement gene expression by qPCR confirmed the RNA-seq finding that SFB gut colonization indeed induced complement expression **(Fig S4C)**. We probed causality by administering AM a cocktail of neutralizing antti-complement antibodies during the first hour in which they were incubated with bacteria.

Antibody-mediated neutralization of complement reduced both uptake and inactivation of bacteria by SFB**^+^** AM, especially SFB**^+^**CA09**^+^** AM, while not impacting these parameters in SFB**^-^** AM. Thus, compared to SFB**^-^** AM, SFB**^+^** AM had elevated complement expression that resulted in an enhanced ability to neutralize bacteria and, moreover, withstand IAV-induced dysfunction **(Fig 3iv and Fig S4E)**.

### SFB colonization systemically enhanced macrophage function while IAV-induced macrophage dysfunction was lung-specific

We next examined the breadth of macrophage dysfunction that resulted from administration of SFB and/or CA09. We found that SFB gut colonization caused enhanced phagocytosis and intracellular bacterial killing by F4/80^+^ macrophages isolated from spleen, small intestine, and heart **(Fig S5A)**. In contrast, CA09-induced macrophage dysfunction was lung-specific. Lung contains two distinct AM populations, namely tissue-resident (TR) and monocyte-derived (Mo) AM. TR-AMs originate from fetal monocytes at birth and can self-renew throughout life, while Mo-AMs differentiate from monocytes recruited to the lung, especially in inflammation. SFB colonization enhanced the antibacterial function of both populations but, in accord with other studies (*15*), only TR-AM displayed dysfunction following CA09 inoculation **(Figure S5B).**

### SFB prevents IAV-induced immune dysfunction

The innate immune response to IAV infection is characterized by high levels of type I interferon (IFN) production, which helps clear the virus but also suppresses the lung’s ability to orchestrate immune responses to bacterial pathogens (*16-19*). SFB colonization profoundly reduced type 1 IFN expression, suggesting SFB**^+^** mice might not exhibit “immune paralysis” following CA09 inoculation. We probed this notion via assaying chemokine (e.g. *s100a9, cxcl3, ccl20, ccl24*) and antimicrobial peptide (*Reg α,b,g*) expression and quantitating innate immune cells in the lung at various times following administration of SFB and/or CA09. As expected, CA09 infection resulted in a robust induction of chemokine expression on day 4 that fully subsided by day 10. *S. pneumoniae*, by itself, also resulted in clear, albeit less broad, lung chemokine expression, which was not observed in CA09/*S. pneumoniae* infected mice, confirming previous observations that IAV induces immune paralysis. SFB colonization did not impact chemokine expression in *S. pneumoniae*-infected mice but alleviated the lack of such chemokine expression in CA09-infected mice. Chemokine expression was mirrored by innate immune cell recruitment. Specifically, *S. pneumoniae* infection resulted in increases in lung neutrophils, eosinophils, and NK cells in SFB**^-^**CA09**^-^** and SFB**^+^**CA09**^+^**, but not SFB**^-^**CA09**^+^**, mice (Fig S6A and Fig S6B). Collectively, these results indicate that SFB colonization ameliorated CA09-induced immune paralysis

### SFB prevented virus-induced immune suppression via impacting AM

We initially suspected that the immune suppression following CA09 infection might reflect the reduction in AM (approximately 50%) caused by the virus. However, a more extensive ablation of AM (about 90%) via clodronate liposomes, by itself, did not prevent *S. pneumoniae*-induced lung chemokine expression. Furthermore, CA09 infection continued to impede *S. pneumoniae*-induced lung chemokine expression in AM-depleted mice **(Fig S6C)**. Thus, the AM depletion was not sufficient or essential for CA09-triggered immune paralysis, although the AM that remained in CA09-infected mice were capable of driving immune paralysis. Specifically, we observed that transplant of AM from CA09-infected mice to AM-depleted mice ablated *S. pneumoniae*-induced lung chemokine secretion and, consequently, neutrophil recruitment. Such immune paralysis was not observed following transfer of SFB**^-^** CA09**^-^**, SFB**^+^**CA09**^-^** or SFB**^+^**CA09**^+^** AM **(Fig 4A)**. Thus, the CA09-induced AM phenotype, rather than CA09-mediated AM ablation, was sufficient for immune paralysis.

**Figure 4.**
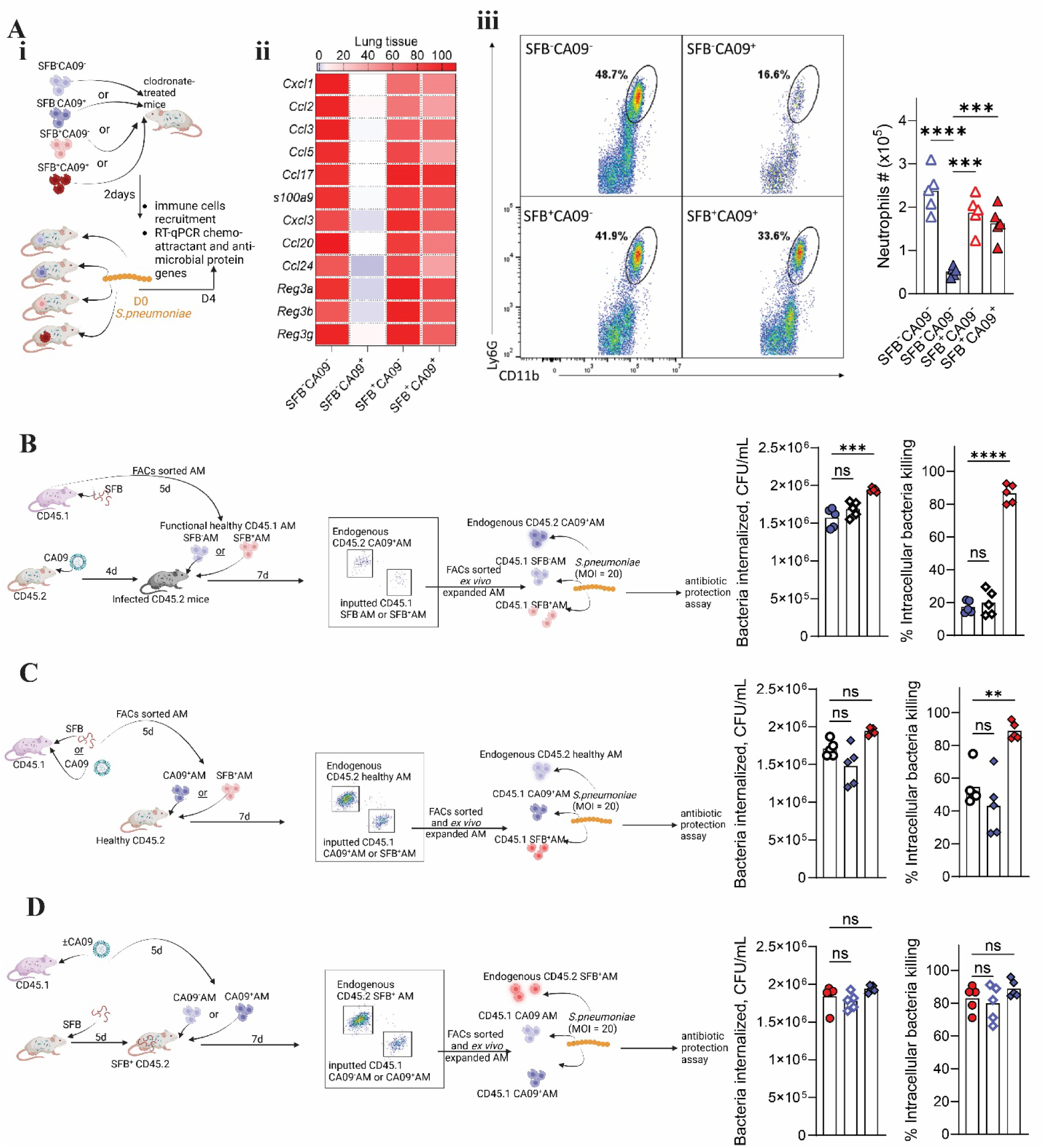
SFB colonization intrinsically reprogrammed AM enabling their high anti-bacterial functions, even in inflamed environments. **(A) (i)** Experimental scheme: SFB^-^mice, colonized with SFB or not 7 days prior were left untreated or inoculated with CA09. Four days post-virus inoculation, 6x10^5^ FACs-sorted SFB^-^ CA09^-^AM, or SFB^-^CA09^+^AM, or SFB^+^CA09^-^AM, or SFB^+^CA09^+^AM were transplanted into clodronate-treated mice. Two days post-transplant, mice were inoculated with *S. pneumoniae* and were euthanized four days later. **(ii)** gene expression of immune cells chemo-attractants and antimicrobial protein of whole lung tissues were analyzed by RT-qPCR and shown as a normalized heatmap. **(iii)** Frequencies and cell numbers of neutrophils were measured by FACs. **(B)** Experimental scheme: SFB^-^AM and SFB^+^AM were FACs-sorted from CD45.1 mice, and 1.5x10^5^ cells were transplanted into CA09-infected mice. Seven days post-transplantation, endogenous CD45.2 CA09^+^AM and transplanted CD45.1 SFB^-^AM and CD45.1 SFB^+^AM were FACs-sorted, and stimulated with live *S. pneumonaie* at MOI = 20, and subjected to an antibiotic protection assay. **(C)** Experimental scheme: CD45.1 mice were orally inoculated with SFB or intranasally inoculated with CA09. Four days later, CD45.1 AM were FACs-sorted and 1.5x10^5^ cells were transplanted into healthy CD45.2 SFB^-^CA09^-^mice. Seven days post-transplantation, endogenous healthy CD45.2 AM, and transplanted CD45.1 CA09^+^AM or SFB^+^AM were FACs-sorted, and subjected to the antibiotic protection assays as in (B). **(D)** CD45.1 mice were inoculated with or without CA09. Four days later, CD45.1 CA09^-^AM or CD45.1 CA09^+^AM were FACs-sorted and 1.5x10^5^ cells were transplanted into CD45.2 SFB^+^ mice. Seven days post-transplantation, endogenous CD45.2 SFB^+^AM and transplanted CD45.1 CA09^-^AM and CA09^+^AM were FACs-sorted and subjected to antibiotic protection assay as in (B). **(B), (C), (D)** Results are shown as CFUL/mL for bacteria internalized in AM, and % of CFU intercellular bacterial killing. n = 5 per group. Results are shown as mean SD. Statistical analysis: One-way ANOVA. ** p < 0.01, *** p < 0.001, **** p < 0.0001, ns not significant.

It appeared likely that a major portion of the difference between SFB**^-^**CA09**^+^** and SFB**^+^**CA09**^+^**, in both influencing lung chemokine expression and managing bacteria they encounter, reflected that the former had been exposed to higher viral loads and, moreover, a highly inflamed type 1 IFN-rich lung environment (*20, 21*). Hence, we examined how SFB**^-^** and SFB**^+^** AM, isolated from CA09**^-^** mice, would fare when transplanted into CA09-infected mice, particularly when transferring small numbers of cells (5x10^4^ cell) to avoid altering of their new environment. SFB**^-^** mice were inoculated with CA09 and administered SFB**^-^** and SFB**^+^** AM 4 dpi. Seven days later, the transplanted cells were re-isolated and assayed for neutralization of an *S. pneumoniae* challenge *ex vivo*. We found that transplanted SFB**^+^** AM retained their high ability to phagocytose and inactivate the pathogen, whereas SFB**^-^**AM matched the dysfunctional phenotype of the endogenous AM **(Fig 4B and Fig S7)**. Conversely, transplant of SFB**^-^**CA09**^+^** AM, which had poor anti-bacterial function, into un-infected mice (i.e. healthy environment) resulted in them acquiring ability to kill *S. pneumoniae* albeit not to the extent of SFB^+^ AM, which maintained their enhanced function in this environment, irrespective of whether they originated from CA09-infected mice **(Fig 4C)**. Lastly, we examined the antibacterial function of SFB**^-^**AM transplanted into lungs of SFB**^+^** mice. We found that both SFB**^-^**CA09**^-^** and SFB**^-^**CA09**^+^** AM fully assumed the ability of the endogenous SFB**^+^**AM to neutralize bacteria **(Fig 4D)**. Thus SFB**^+^** AM hold their phenotype in various environments while SFB**^-^** AM tend to assume the AM phenotype that predominates their environment.

### Downregulation of AM Notch4 on AM protects against CA09-induced dysfunction

The extent to which SFB colonization of the intestine, by itself, impacts AM gene expression is fairly modest. Specifically, analysis of AM gene expression by RNA-seq found that SFB gut colonization induced expression of only a couple dozen genes and reduced expression of but one, namely Notch 4 **(Fig S4C)** (*8*). We previously observed that antibody-mediated blockade of Notch 4 on AM could partially recapitulate the impact of SFB colonization via assaying AM *ex vivo* or following their transplant into mice (*11*). Hence, we deployed these approaches here to examine the role of Notch 4 CA09-induced AM dysfunction. SFB**^+^** and SFB**^-^** AM, treated with anti-Notch 4 or isotype control antibody, were exposed to UV-inactivated IAV and then challenged *ex vivo* or *in vivo* (i.e. following transplant into SFB-mice) with *S. pneumoniae*. Anti-Notch treatment improved AM antibacterial function *ex vivo* and *in vivo*, wherein recipients of anti-Notch 4-treated AM exhibited greater chemokine expression, neutrophil levels and lower lung CFU, albeit less than recipients of SFB**^+^** AM **(Figure 5A)**. The increased antibacterial function of anti-Notch 4-treated and SFB**^+^** AM, relative to SFB**^-^** AM, were paralleled by reduced levels of UV-CA09-induced Type I IFN expression **(Fig S8A)**. Blockade of Notch 4 *in vivo*, via intraperitoneal administration of the antibody, immediately prior to inoculation with CA09 resulted in endogenous AM displaying high antibacterial functionality and, moreover, a lung environment into which SFB-AM would have similar functionality albeit not quite the level exhibited by SFB**^+^** AM **(Fig 5B)**. In contrast, administration of anti-Notch 4-treated SFB**^-^** AM into an inflamed IFN-rich environment, i.e. lungs of CA09-infected SFB**^-^** mice, resulted in such AM acquiring the dysfunctional phenotype of the endogenous AM while SFB^+^ AM maintained high antibacterial function in this environment **(Fig 5C)**. Such ability of Notch 4 blockade to impact CA09-induced IFN but not benefit AM once an IFN-rich environment was established correlated with the time course of CA09-induced Notch 4 expression and AM dysfunction **(Fig S8B)**. Thus, the down regulation of AM Notch 4 expression following SFB colonization contributes to early events that help SFB^+^ AM maintain antibacterial function but other mechanisms allow SFB**^+^** AM to maintain such functions.

**Figure 5.**
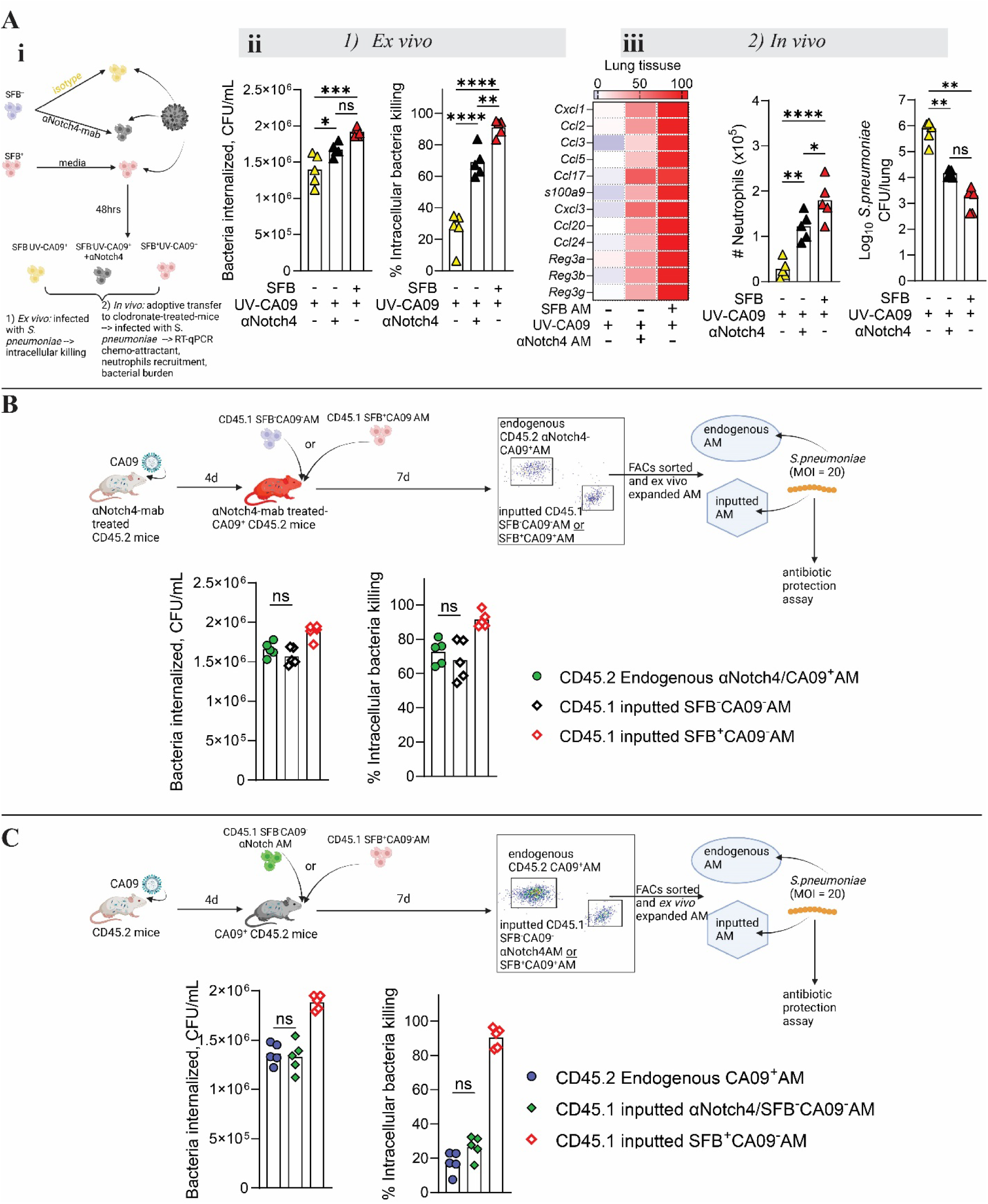
SFB’s down-regulation of Notch4 played a role in initiating, but not sustaining, AM antibacterial functions amidst IAV infection. **(A) (i)** Experimental scheme: SFB^+^AM and SFB^-^AM were exposed to inactivated UV-CA09 with Notch4 neutralizing or isotype control antibodies. **(ii)** *Ex vivo* experiments: 48 hours later, AM were challenged with live *S. pneumoniae* and subjected to antibiotic protection assay. Results shown as Results are shown as CFUL/mL for bacteria internalized in AM, and % of CFU intercellular bacterial killing. **(iii)** *In vivo* experiment: 6x10^5^ SFB^-^UV-CA09^+^AM, or SFB^-^UV-CA09^+^AM/αNotch4 AM, or SFB^+^UV-CA09^+^AM were transplanted into clodronate-treated mice and challenged with *S. pneumonaie*. Four days later, RNA was harvested from whole lung tissues, gene expression was assayed by RT-qPCR, and results were normalized and displayed as a heatmap, neutrophil infiltration was analyzed by FACs, and lung bacterial burden weas assayed by counting CFU on blood agar plate. **(B)** Experiment scheme: αNotch4 mAB-treated CD45.2 mice were inoculated with CA09, and four days post-inoculation, mice were transplanted with 1.5x10^5^ CD45.1 SFB^-^CA09^-^AM or CD45.1 SFB^+^CA09^-^AM. Seven days post transplantation, endogenous CD45.2 αNotch4/CA09^+^AM and transplanted AM were FACs-sorted, and challenged with live *S. pneumoniae* at MOI = 20, and then subjected to an antibiotic protection assay. (C) Experimental scheme: CD45.2 mice were inoculated with CA09, and four days later, mice were transplanted with 1.5x10^5^ CD45.1 αNotch4/SFB^-^CA09^-^AM or CD45.1 SFB^+^CA09^-^ AM. Seven days post transplantation, endogenous CD45.2 CA09^+^AM and transplanted AM were FACs-sorted, stimulated with *S. pneumoniae* at MOI = 20, and then subjected to the antibiotic protection assay. **(B), (C)** Results are shown as CFUL/mL for bacteria internalized in AM, and % of CFU intercellular bacterial killing. All experiments were performed with n = 5 per group. Results are shown as mean SD. Statistical analysis: * p < 0.05, ** p < 0.01, *** p < 0.001, **** p < 0.0001, ns not significant.

### Adoptive transplant of SFB^+^AM mitigated the severity of secondary bacterial infections

Lastly, we performed AM transplantation studies to assess the extent to which the changes in AM, induced by SFB gut colonization, were sufficient to protect against CA09/*S. pneumoniae* infection. First, we utilized an approach in which transplanted AM would be present throughout CA09 and *S. pneumoniae* infection **(Fig 6A)**. Specifically, SFB**^-^** mice were subjected to clodronate-mediated AM depletion and then administered SFB**^-^** or SFB**^+^** AM. Such mice were then inoculated with CA09 and, 4 days later, *S. pneumoniae*. Recipients of SFB**^-^** and SFB**^+^** AM displayed starkly different outcomes following this regimen, namely 0 and 100% survival, respectively, which correlated with a three orders of magnitude difference in *S. pneumoniae* CFU and marked differences in gross and histopathologic lung appearance 4 days post-inoculation. A significant portion of this difference likely reflected differential responsiveness of these groups of mice to CA09. Indeed, we have previously shown that SFB**^+^** AM resist depletion following transplant and, concomitantly, their recipients are relatively resistant to CA09 infection as assessed by viral titers and clinical signs. Accordingly, we herein observed recipients of SFB**^-^** AM had lost more weight indicating more pronounced disease and, concomitantly, had far fewer AM at the time when they were exposed to *S. pneumoniae.* Thus, we adjusted our experimental scheme to assess the potential impact of SFB**^+^** AM on secondary bacterial infection that were not a direct consequence of their effect on CA09 infection **(Fig 6B)**. Specifically, SFB**^-^** mice were infected with CA09, administered SFB**^-^** or SFB^+^ AM 6 dpi with CA09, and then, 2 days later, inoculated with *S. pneumoniae*. Compared to mice receiving SFB**^-^** AM, recipients of SFB**^+^** AM displayed 3-fold lower levels of lung CFU 4 days after *S. pneumoniae* inoculation.

**Figure 6.**
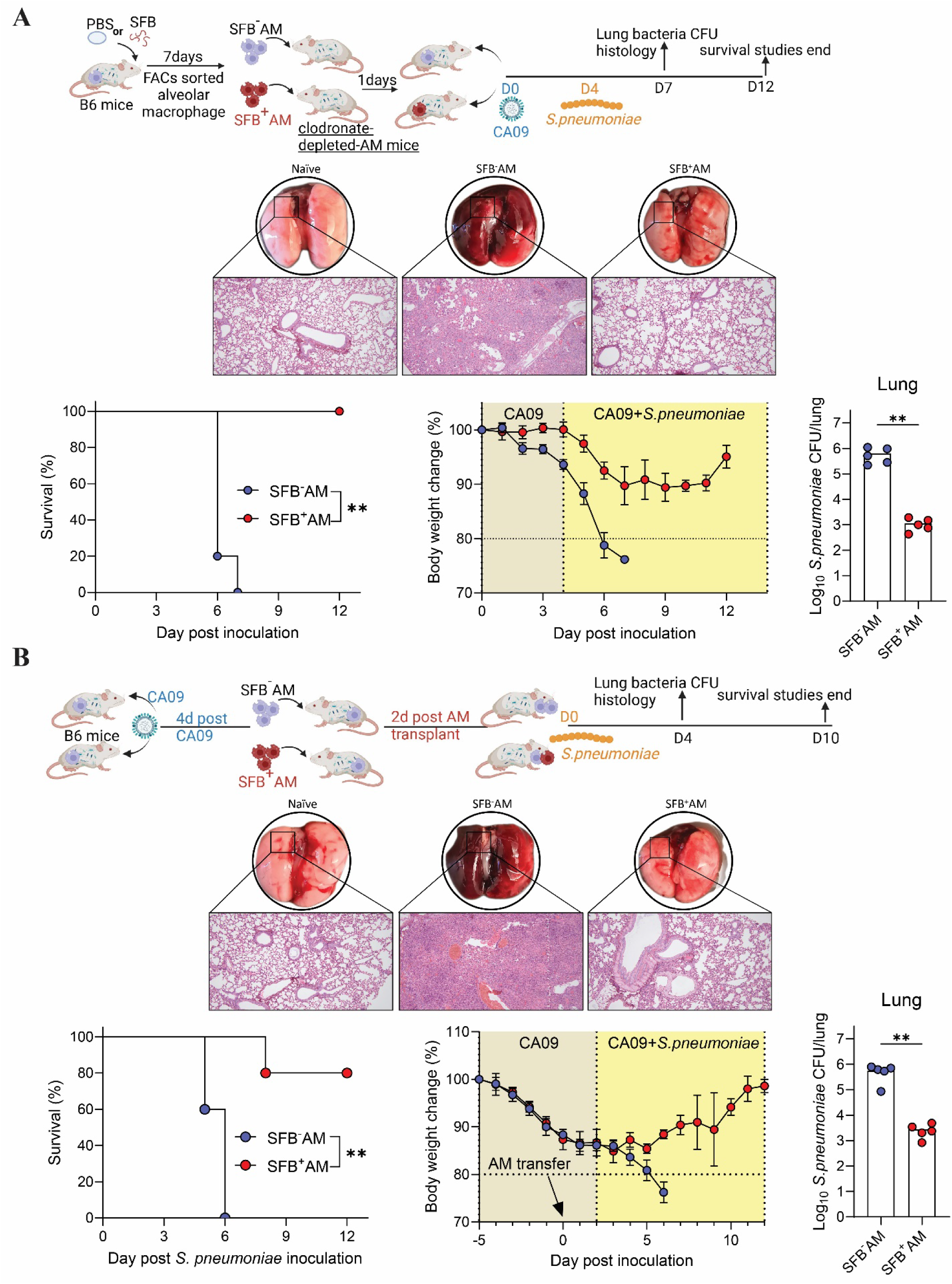
Transplant SFB^+^AM conferred resistance to secondary bacterial infection. **(A)** As schematized, SFB^-^mice were administered SFB or PBS. Seven day later, AM were FACs-sorted and 6x10^5^ cells were transplanted to a clodronate-mediated-AM-depletion-SFB^-^mice. One day post-transplant, mice were inoculated with 50 TCID_50_ CA09 and four days later with 1x10^7^ *S. pneumoniae*. **(B)** As schematized, SFB^-^mice were inoculated with 50 TCID_50_ CA09. On day four post CA09 inoculation, mice received 6x10^5^ AM FACs-sorted from SFB^+^ or SFB^-^ mice. **(A) and (B)** Two days post AM transfer, mice were inoculated with 1x10^7^ *S. pneumoniae.* Mice were euthanized four days post bacteria inoculation for lung bacteria titer, gross lung and histology assessment or monitored for survival and body weight. n = 5 per group. Data are representative of two independent experiments, yielding an identical pattern of results. Results are shown as mean SD. Statistical analysis: lung bacteria burden: student’s t test. Survival: log-rank Mental-Cox Test. *p < 0.05, ** p < 0.01.

Moreover, recipients of SFB**^-^** AM uniformly succumbed within 6 days of the secondary *S. pneumoniae* challenge, whereas most (80%) of recipients of SFB**^+^** AM survived and regained the weight lost in response to the primary CA09 infection. Thus, reprogramming of AM by gut SFB colonization not only maintains the presence of AM following respiratory viral infection but confers those cells with enhanced ability to manage respiratory bacterial pathogens.

## DISCUSSION

We herein report that one gut microbiota constituent, SFB, provided stark protection in mouse models of secondary bacterial infection following IAV infection. Such protection was largely mediated by AM, whose reprogramming by SFB colonization conferred these cells with ability to maintain high functionality amidst polymicrobial onslaught. This finding was not entirely surprising in that we have previously shown that SFB-colonized mice resist IAV infection and AM depletion, which had been implicated as contributing to the high proneness to respiratory bacterial pathogens following IAV infection (*10*). Indeed, a portion of the protection against secondary bacterial infection conferred by SFB likely reflected that SFB-colonized mice developed less severe disease and had more AM at the time we exposed mice to bacterial pathogens. Nonetheless, given the high disease burden caused by secondary bacterial infections, including those studied here, namely *S. pneumoniae, S. aureus, and H. influenzae*, in driving complicated viral disease, the understanding that the presence of one additional microbiota constituent protected against these challenges is an important advance. Furthermore, irrespective of AM numbers, SFB-induced alterations in AM phenotype played a key role in mediating the ability of SFB-colonized mice to resist lung bacterial pathogens.

AM from SFB colonized mice were, on a per cell basis, intrinsically more adept at managing bacterial pathogens even prior to IAV infection. These findings mirror our recent report that such SFB-induced AM alterations confer greater ability to neutralize IAV. Furthermore, similar mechanisms were involved in that the enhanced ability of AM from SFB-colonized mice to neutralize bacterial pathogens and IAV involved enhanced phagocytosis and complement production. On the one hand, the well appreciated role of complement in mediating phagocytosis and neutralization of bacteria is consistent with our current observation of greater bacterial clearance by SFB**^+^** AM. On the other hand, our results do not follow the long held general paradigm that innate immune cells are generally in either an antiviral state or antibacterial state such that they are seldom optimized to handle both classes of pathogens.

The level of enhanced antibacterial immunity conferred by SFB was relatively modest by itself but became dramatic following IAV infection, even when disregarding that SFB-colonized mice resisted AM depletion. Indeed, IAV infection starkly impaired the ability of SFB**^-^** AM to neutralize internalized bacteria but only slightly decreased this function in SFB**^+^** AM. A portion of this differential impact likely reflected that reductions in AM anti-bacterial potency were mediated by type I IFN expression of which is dramatically reduced in lungs of SFB**^+^** mice following IAV infection. Reduced IFN expression in SFB**^+^** mice is mediated, in part, by SFB- induced suppression of Notch 4. Here, we observed that blockade of Notch 4 reduced IAV-induced AM IFN expression and, likely as a consequence, initiation of dysfunction, but did reduce their dysfunction when such AM were placed into an inflamed IFN-rich environment. In contrast, even when placed into an inflamed IFN-rich environment, SFB**^+^** AM maintained a high level of anti-bacterial function, arguing that SFB reprogramming imbued them with the wherewithal to maintain core tenets of innate immune function irrespective of their environment. We hypothesize a role for SFB-induced chromatin remodeling in mediating such stability of the SFB**^+^** AM phenotype.

Enhanced bacterial neutralization by SFB**^+^** AM, combined with these cells having withstood IAV-induced depletion and being more abundant, were likely major contributors to the reduced bacterial loads in the lungs of SFB-colonized mice. In turn, such reduced bacterial loads correlated with, and highly likely contributed to, the symptoms of secondary bacterial infection (i.e. weight loss, pneumonia, and death). Additionally, SFB-colonized mice retained the ability to express chemokine genes and, consequently, orchestrate recruitment of other innate immune cells in response to secondary bacterial infection. That AM ablation via clodronate did not reduce this ability argues that it likely reflects epithelial cell orchestration of innate immunity and, in any case, does not require AM per se. Rather, AM transplant studies indicated that SFB**^-^**CA09**^+^** blocked lung recruitment of immune cells while SFB**^+^**CA09**^+^** AM did not impede it. IAV-induced suppression of immune cell movement was likely mediated by type I IFN, which was expressed in response to IAV by SFB**^-^** but not SFB**^+^** AM. That SFB**^+^** mice retain the ability to recruit immune cells likely aids in withstanding bacterial challenges.

In conclusion, SFB-induced reprogramming of AM, which we previously showed reduces severity of RVI, also enhances antibacterial innate immunity. The impact of the latter is especially pronounced in post-RVI secondary bacterial infections. Such infections are major contributors to complicated RVI disease and poor outcomes. Thus, we speculate that harnessing the mechanism by which SFB colonization reprograms AM may provide a means of mitigating disease burden associated with RVI.

## METHODS

### Mice

Wild-type C57BL/6 (B6 WT) mice were purchased from The Jackson Laboratory and used at 3 weeks of age. Experiments were conducted using age- and gender-matched groups. The Jackson Laboratory does not routinely test for segmented filamentous bacteria (SFB) in mice; however, all mice obtained from this vendor were tested and confirmed to be SFB negative upon arrival at GSU. Animal studies were approved by the Institutional Animal Care and Use Committee (IACUC) of Georgia State University.

### Mono-associated SFB transplantation

Fecal samples were collected from germ-free donor mice, suspended in 20% glycerol PBS solution at 80 mg/mL concentration, passed through a 40 µm filter, and stored at -80°C. Frozen fecal suspensions were orally administered to recipient mice at 200 ml per mouse.

### Virus and Bacteria strains

*Streptococcus pneumoniae* 6303 (strain designation CIP 104225), Methicillin-resistant *Staphylococcus aureus* (strain designation F-182), was obtained from the American Type Culture Collection (ATCC). Frozen bacterial stocks of S. pneumoniae were streaked out on Trypticase Soy Agar plates containing 5% sheep blood. A single colony was inoculated into 5 ml of Todd-Hewitt broth with 0.5% added yeast extract and grown at 37°C with 5% CO_2_ without shaking. *Haemophilus influenzae* (strain designation AMC 36-A-5 624, NCTC 8469) was obtained from ATCC and cultured in ATCC medium 2167 Haemophilus Test Medium media 37°C with 5% CO_2_ for 48 hours. Bacteria were centrifuged and resuspended in PBS and diluted to the appropriate intended inoculum (1x10^7^ CFU or 500 CFU per mouse). Recombinant A/CA/07/2009 (H1N1) (CA09) GFP-expressing reporter virus version of A/CA/07/2009 (H1N1) was generated as previously described

### Virus and bacterial co-infection

For virus infections, animals were anesthetized with isoflurane and intranasally inoculated with 50µl of 50 TCID_50_ units/animal of recombinant A/CA/07/2009 (H1N1) (CA09) GFP-expressing reporter virus. For secondary or co-bacterial infections, anesthetized mice were intranasally inoculated with 50 µL of PBS containing 1x10^7^ CFU of *S. pneumoniae*, or *S. aureus*, or *H. influenzae*. Bacterial burdens in the lungs, BALF, and spleen were measured by euthanizing the infected mice on either day 4 or day 5 post-infection and plating serial 10-fold dilutions of each sample onto blood agar plates (*S. pneumoniae* and *S. aureus*) or chocolate agar plates (*H. influenzae*).

### Lung digestion and flow cytometry analysis of immune cells

Lungs were dissected from mice and minced with scissors before digestion in RPMI solution containing 5% FBS, DNase I, and 1 mg/mL Collagenase type IV (Sigma) for 30 minutes at 37°C. During digestion, lungs were further homogenized by swishing with a 5 mL syringe every 10 minutes. For flow cytometry analysis, cells were blocked with 1 mg/million cells using anti-CD16/anti-CD32 in 100 µL PBS for 15 minutes at 4°C, followed by washing with PBS to remove any residual blocking antibodies. Cells were then incubated with the following conjugated monoclonal antibodies: CD45 BV605, MHCII BV650, CD11b APC-Cy7, CD11c BV786, Ly6C PerCp-Cy5.5, Ly6G AF700, CD64 BV421, CD24 FITC, CD117 GV510, SiglecF PE-CF594, and Fcer1 APC. Multi-parameter analysis was performed on a CytoFlex (Beckman Coulter) and data were analyzed using FlowJo software (Tree Star). The gating strategy for innate immune cells was based on previous described (*11, 22*).

### Antibiotic protection/Intracellular bactericidal assay

Alveolar macrophages were sorted, cultured, and incubated with *S. pneumoniae*, or *S. aureus*, or *H. influenzae* at a multiplicity of infection (MOI) of 20. The cultures were centrifuged at 500 x g for 10 minutes to increase macrophage-bacteria interaction, followed by a one-hour incubation for bacterial uptake. The media was then replaced with an antibiotic cocktail (RPMI with 5% FBS, 500 µg/mL gentamicin, 500 U/mL penicillin, and 500 µg/mL streptomycin) and incubated for 1 hour to eliminate extracellular bacteria. After washing with PBS, some wells were lysed with 0.1% Triton X-100 to collect bacterial CFUs, which were serially diluted and plated on blood agar plates or chocolate agar plates to measure bacterial uptake. The remaining wells were incubated for an additional hour in fresh media before lysis and CFU collection to assess bacterial survival. The percentage of CFU remaining was calculated as 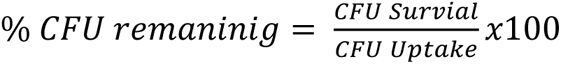

### Histopathologic analysis

Lungs were perfused using 10% neutral-buffered formalin, dissected, and fixed for X days. Formalin-fixed lungs were transferred to 70% EtOH, embedded in paraffin, sectioned, stained.

### Adoptive transplant of alveolar macrophages

Lungs were harvested from C57BL/6 mice and digested with Collagenase type IV as described previously (ref). Alveolar macrophages (defined as CD45^+^MHCII^+^CD11c^+^SiglecF^+^) were stained and FACS-sorted. Depend on experiment either 5x10^4^ or 6x10^5^ alveolar macrophages in 30 µL PBS were administered intranasally to recipient mice.

### RNA-seq analysis

RNA-seq analysis were performed with data sets from (ref CHM paper). Total lung RNA-seq accession number: PRJEB71449 and alveolar macrophages RNA-seq accession number: PRJEB71689 from ENA European Nucleotide Archive. Pathway enrichment analysis was analyzed using GSEA software.

### *Ex vivo* inhibiting Notch 4 signaling in alveolar macrophages

Alveolar macrophages were FACS-sorted as described (ref). After sorting, cells were treated with either isotype control antibody (Cat# BE0091, BioXCell) or an anti-Notch4 monoclonal antibody (10mg/mL) (Cat# BE0129, BioXCell), known to neutralize Notch 4 for 45 minutes.

### Alveolar macrophage phagocytosis assay

AM were FACS-sorted and 1x10^5^ were seeded in 96 well incubated overnight. Cells were washed and incubated with pHrodo S. aureus BioParticle for 2 hours. Level of cell fluorescence, an indicator of bioparticle uptake, was measured via flow cytometry.

**Table 1:**
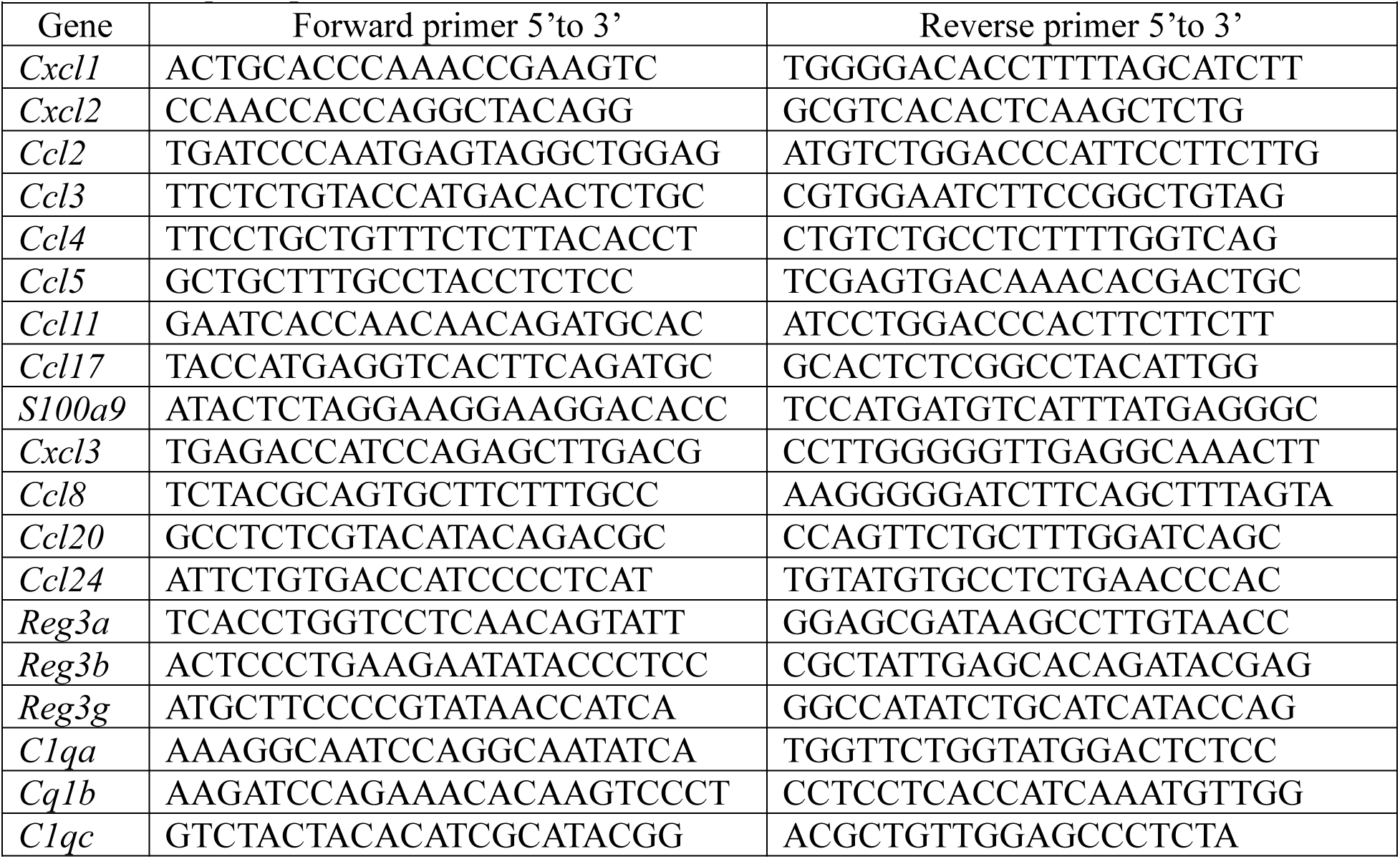
RT-qPCR primers.

### Quantitative real-time PCR

Total RNA from lungs or from FACs-sorted alveolar macrophages were isolated using the Qiagen RNeasy Mini Kit with on-column DNase digestion according to manufacturer’s protocol. cDNA was generated using the Superscript First Strand Synthesis kit for RT-PCR and random hexamer primers (Invitrogen). RT-qPCR was performed with SYBR Green using StepOnePlus PCR system (Applied Biosystem), and gene expression was normalized to *Gapdh*. All primers are listed below.

### Quantification and statistical analysis

Results were expressed as mean ± SEM. All data was plotted in GraphPad Prism version 10. Statistical significance was assessed by One-way ANOVA, Student’s t test, and Two-way ANOVA. Differences between experimental groups were considered significant if *p<0.005, **p<0.005, ***p<0.001, ****p<0.0001.

### Experimental schematics and graphical abstract

All experimental schematics and the graphical abstract were created using BioRender (https://biorender.com/).

---

**Figure S1.**
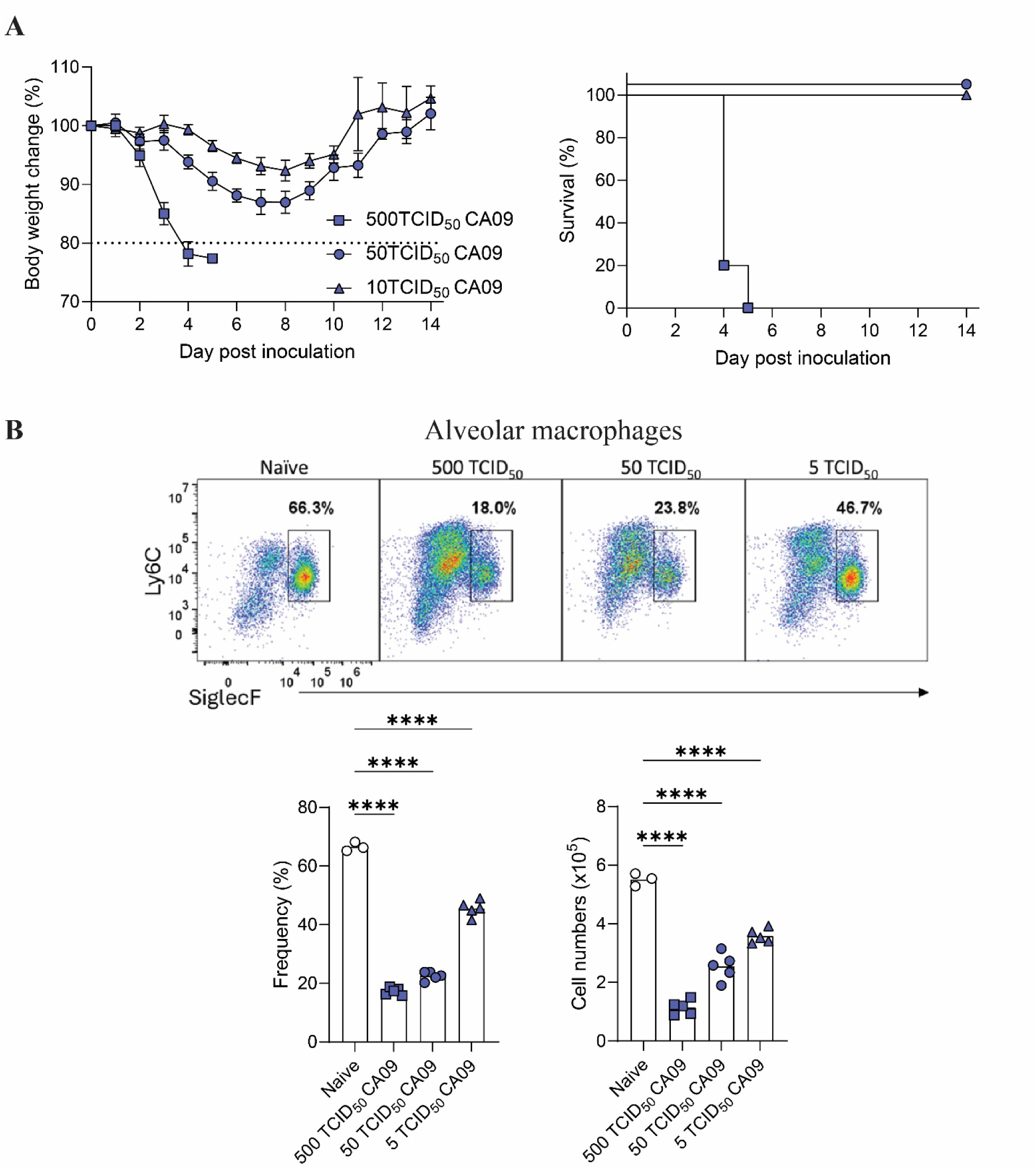
Defining a non-lethal influenza virus (CA09) dose to study secondary bacterial infection. **(A)** Three-week-old C57BL/6 mice were inoculated with the A/CA/07/2009 (H1N1) influenza virus (CA09) at different doses: 500 TCID_50_, 50 TCID_50_, and 5 TCID_50_. Survival and body weight were monitored for fourteen days. **(B)** A representative flow plot and the frequency and cell numbers of alveolar macrophages (AM) were analyzed on day 4.5 post-CA09 inoculation. All experiments n = 3-5 per group. Results are shown as mean SD. Statistical analysis: One-way ANOVA. ****p < 0.0001.

**Figure S2.**
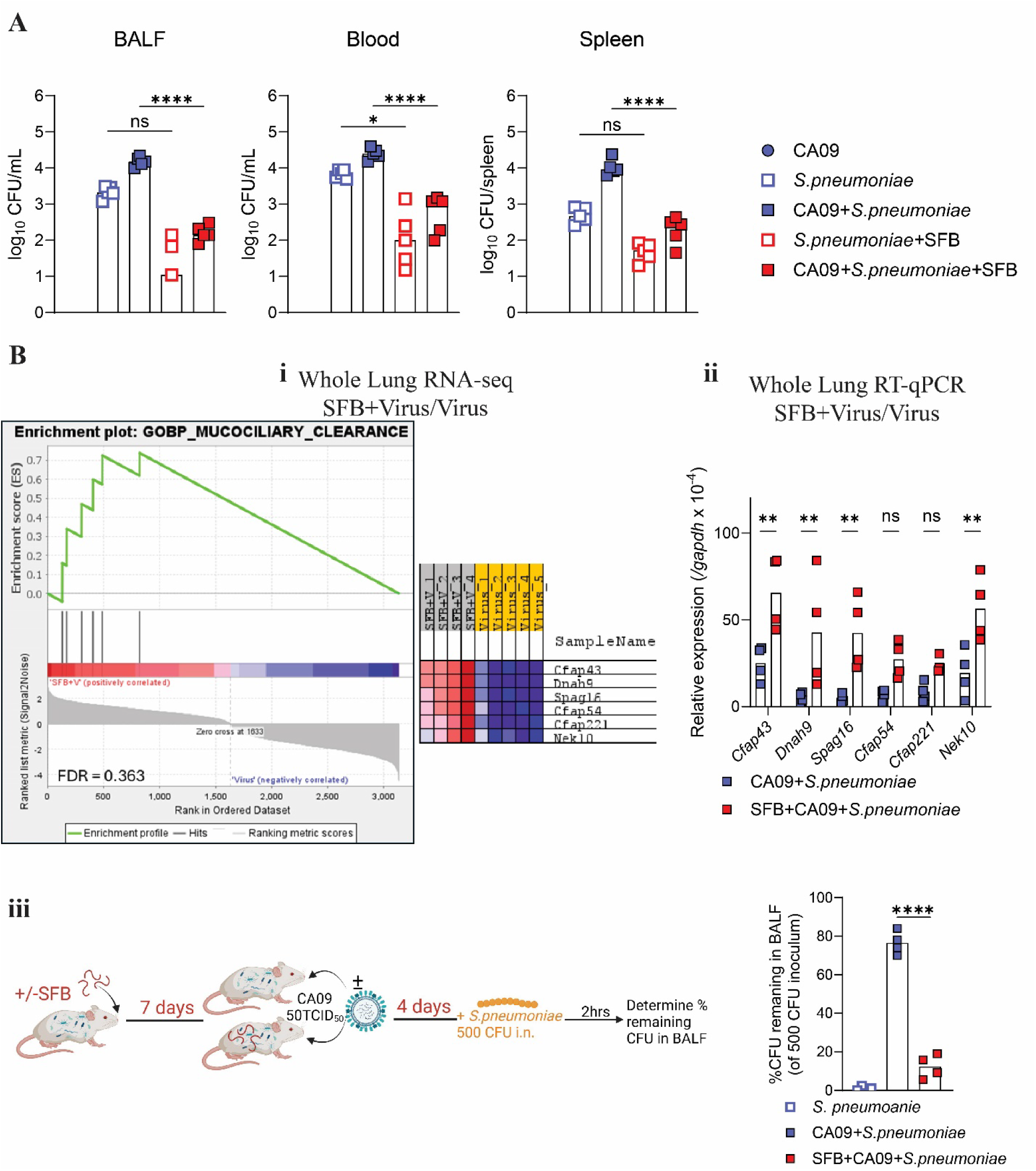
Resistance of SFB-colonized mice to secondary/co-bacterial infection associated with enhanced mucociliary clearance. **(A)** Bacteria titer in bronchoalveolar fluid (BALF), blood and spleen of mice in figure 2A. **(B) (i)** Enrichment plot from gene set of mucociliary clearance calculated by GSEA analysis and **(ii)** RT-qPCR of genes relate to mucociliary clearance from total lung RNA-seq of SFB+CA09/CA09. **(iii)** Three-week-old C57BL/6 SFB^-^ mice were either colonized with SFB or not and inoculated with CA09. Four days post-virus inoculation, the mice were inoculated with intranasally 500 CFU *S. pneumoniae*. Two hours after bacterial inoculation, BALF harvest bacterial remaining CFU in BALF was calculated. % remaining CFU in BALF = 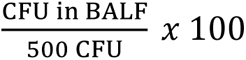. All experiments n = 5 per group. Data are representative of two experiments, yielding an identical pattern of results. Results are shown as mean SD. Statistical analysis: One-way ANOVA. *p < 0.05, ** p< 0.01, ****p < 0.0001, ns not significant.

**Figure S3.**
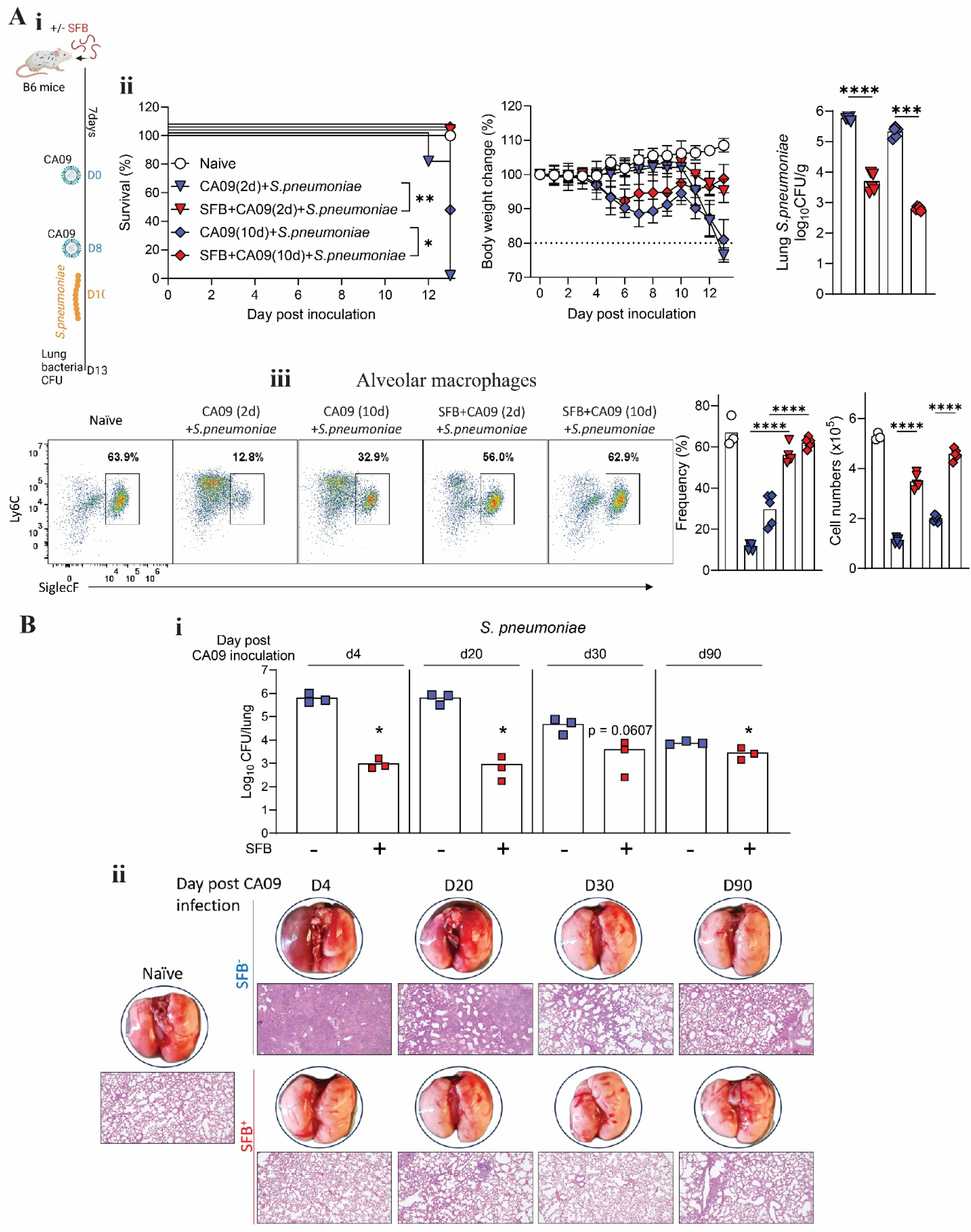
SFB-mediation protection against secondary bacterial infection persisted for several weeks. **(A) (i)** Experimental scheme: Three-week-old C57BL/6 SFB^-^ mice were either colonized with SFB or not and inoculated with CA09. At two days, or ten days post-virus inoculation, mice were inoculated with intranasally with 1x10^7^ CFU *S. pneumoniae.* Mice were either monitored for body weight loss and survival rate or euthanized on day four post bacterial inoculation. **(ii)** Survival rate, body weight, and lung bacterial titer. **(iii)** Representative flow plot, frequency, and cell numbers of alveolar macrophage. **(B)** SFB^-^ mice were either colonized with SFB or not and inoculated with CA09. At four days, twenty days, thirty days or ninety days post-virus inoculation, mice were inoculated with intranasally with 1x10^7^ CFU *S. pneumoniae* and euthanized four days later. Lung bacterial burden and lung histology were assessed. All experiments performed with n = 3-5 per group. Results are shown as mean ± SD. Statistical analysis: lung bacterial burden: One-way ANOVA or Student’s t test. Survival: log-rank Mental-Cox Test. *p < 0.05, ***p < 0.001, ****p < 0.0001.

**Figure S4.**
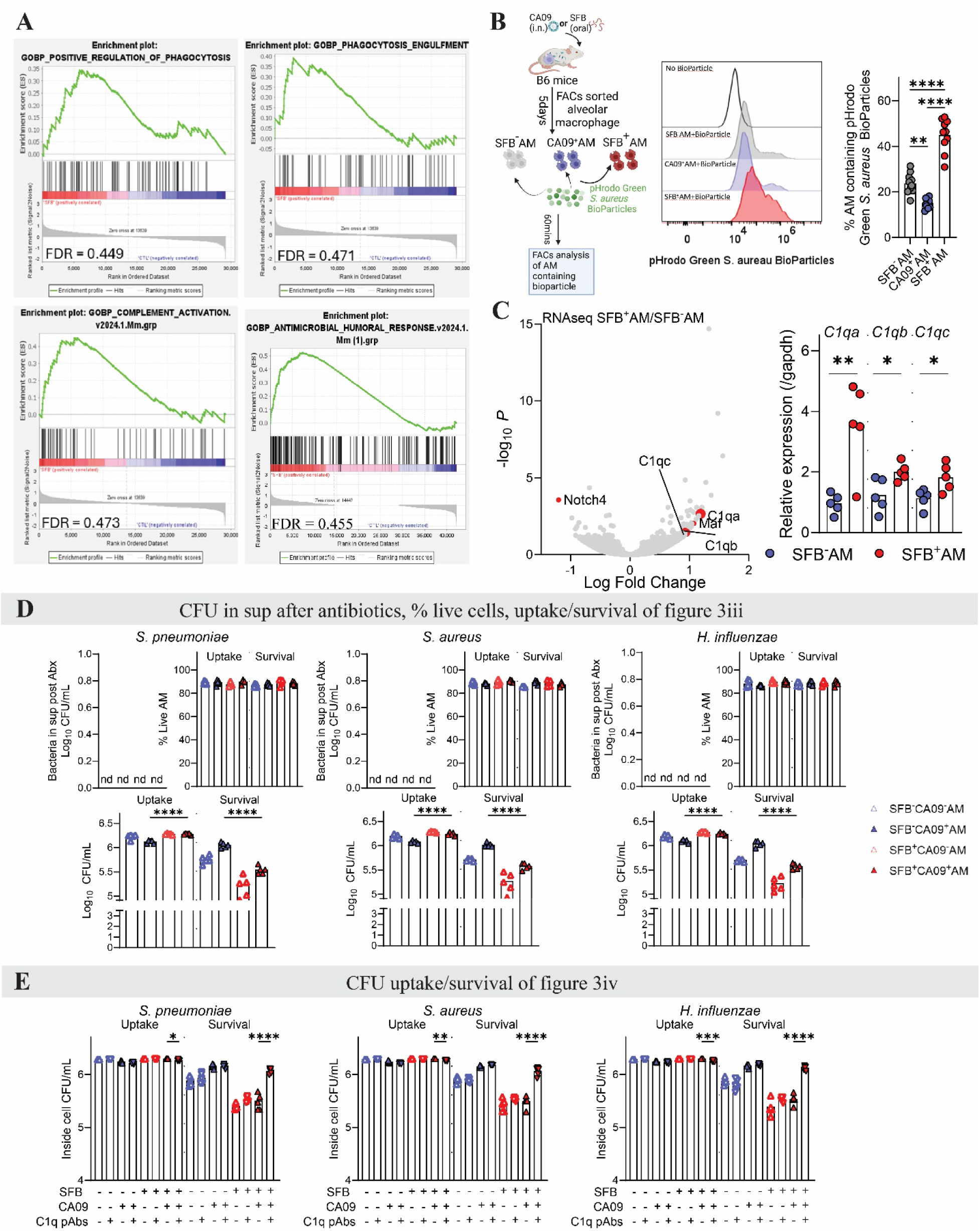
Increased phagocytosis and complement activation in SFB^+^AM. **(A)** Enrichment plot from gene set of positive regulation of phagocytosis, phagocytosis engulfment, complement activation, and antimicrobial humoral responses calculated by GSEA analysis from RNA-seq of SFB^+^AM/SFB^-^AM. **(B)** C57BL/6 SFB^-^ mice were either colonized with SFB or inoculated with CA09 or received PBS for five days and AM were FACs-sorted and exposed to pHrodo Green *S. aureus* BioParticle for 30mins. Degree of phagocytosis were assayed by FACs. **(C)** The volcano blot of RNA-seq and RT-qPCR shows C1qa, C1qb and C1qc are up-regulated in SFB^+^AM. **(D)** log_10_CFU counts of Figure 3iii. Bacterial in supernatants post antibiotics, % live macrophage, CFU/mL count of Uptake and Survival of bacteria inside AM. **(E)** CFU/mL count of Uptake and Survival of bacteria inside AM of Figure 4iv. All experiments n = 3-5 per condition. Results are shown as mean ± SD. Statistical analysis: One-way ANOVA. *p < 0.05, **p < 0.01, ***p < 0.001, ****p < 0.0001, ns not significant.

**Figure S5.**
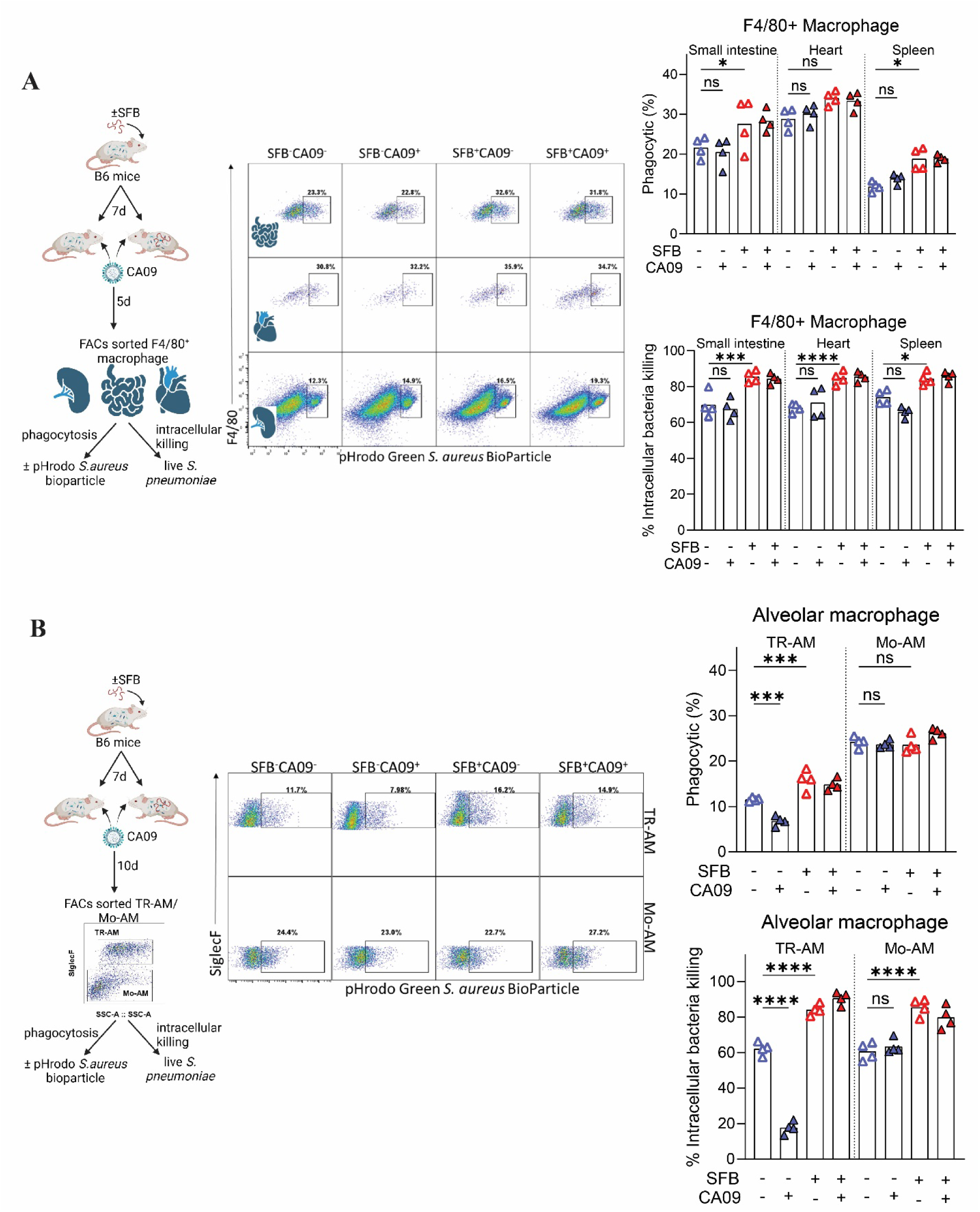
SFB systemically enhanced macrophage anti-bacterial function while CA09- induced dysfunction was restricted to the lung-resident macrophages. **(A)** Experimental scheme: SFB^-^ mice were colonized with SFB, and seven days post-colonization, mice were inoculated with CA09. F4/80^+^ macrophages from the spleen, small intestine, and heart were FACS-sorted and either stimulated with pHrodo Green *S. aureus* BioParticles to measure phagocytosis or with live *S. pneumoniae* to assess intracellular killing abilities. FACS plots, percentages of phagocytic cells, and percentages of remaining bacteria inside macrophages were shown. **(B)** SFB^-^ mice were colonized with SFB and inoculated with CA09 as described in (A). Ten days post-CA09 inoculation, Mo-AM and TR-AM were FACS-sorted and exposed to pHrodo Green *S. aureus* BioParticles or live *S. pneumoniae*. FACS plots, percentages of phagocytic cells, and percentages of intracellular killing of AM were shown. All experiments were performed with n = 4 per group. Results are shown as mean ± SD. Statistical analysis: *p < 0.05, ***p < 0.001, ****p < 0.0001, ns: not significant.

**Figure S6.**
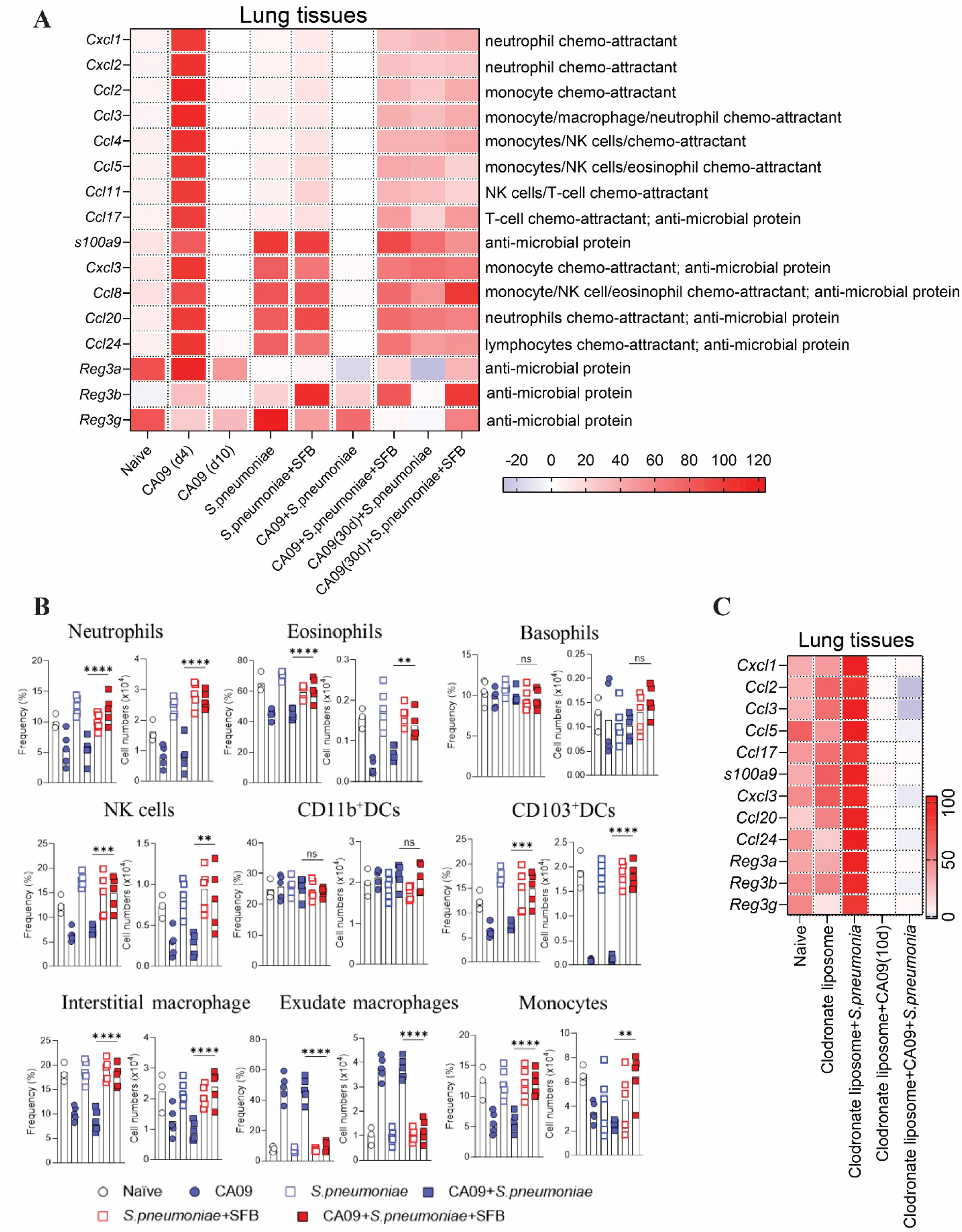
SFB prevented influenza virus-induced blockade in *S. pneumoniae*-induced immune cell recruitment. **(A)** SFB^-^mice were colonized, or not, with SFB, 7 days later, inoculated with CA09. Fours, or ten or thirty-days post-virus inoculation, mice were intranasally inoculated with 1x10^7^ CFU of *S. pneumoniae*. Four days post bacterial challenge, RNA was extracted from whole lungs, and gene expression was analyzed by RT-qPCR and normalize genes expression shown as a heat map. **(B)** Frequencies and cell numbers of innate immune cells were assessed by FACs. **(C)** Mice were treated with clodronate liposomes or control liposomes and inoculated with *S. pneumoniae* alone or co-infected with CA09/*S. pneumoniae*. Four days post-inoculation, RNA was extracted from whole lungs, and gene expression was analyzed by RT-qPCR and results were normalized and shown as a heatmap. All experiments were performed with n = 3-5 per group. Results are shown as mean ± SD. Statistical analysis: *p < 0.05, **p < 0.01, ***p < 0.001, ****p < 0.0001.

**Figure S7.**
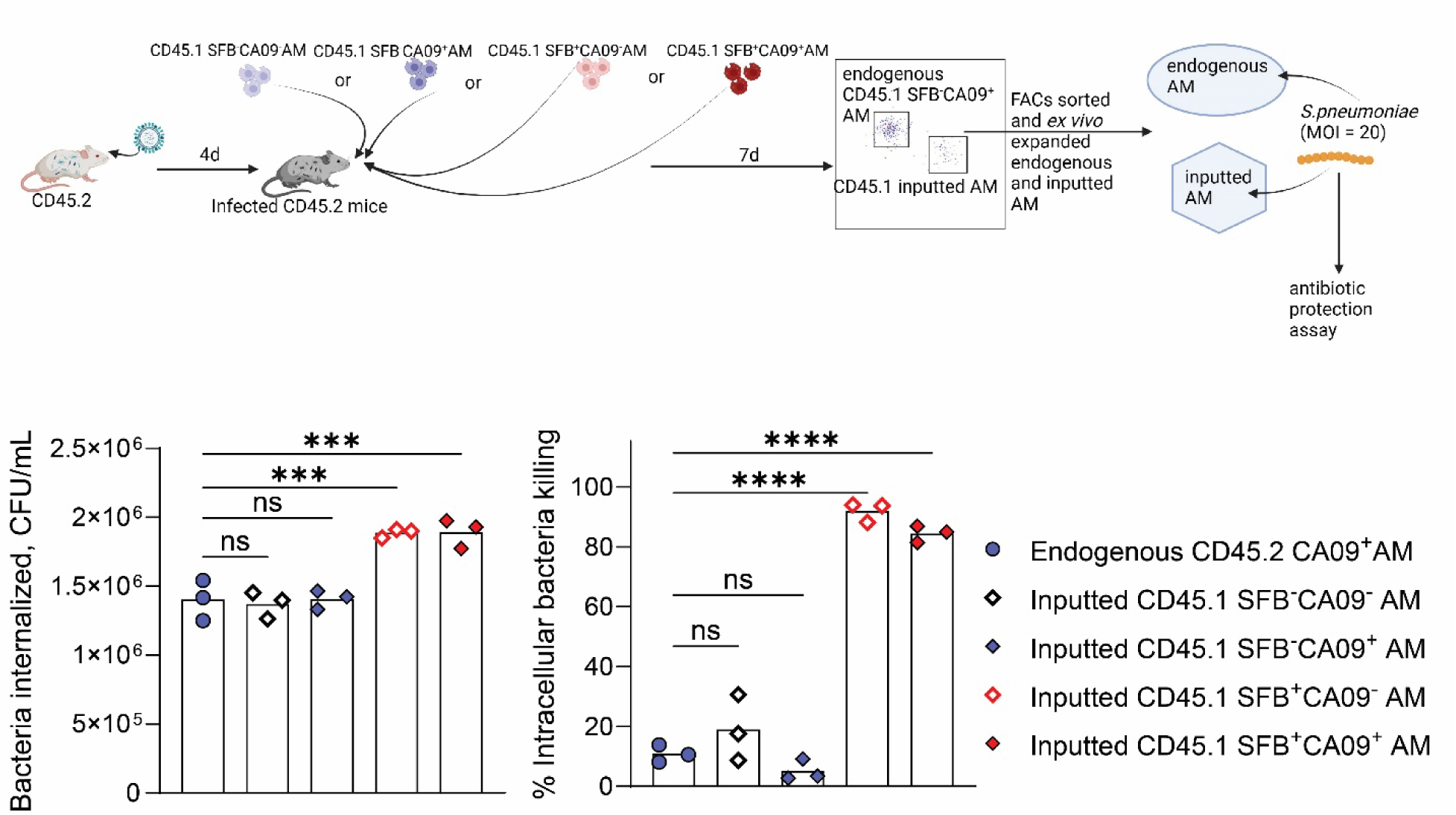
SFB-reprogrammed AM retained antibacterial function following transplant into in IAV lungs. As shown in the schematic, CD45.2 mice were inoculated with CA09, and four days later, transplanted with 1.5x10^5^ CD45.1 SFB^-^CA09^-^AM, or SFB^-^CA09^+^ AM, or CD45.1 SFB^+^CA09^-^ AM, or CD45.1 SFB^+^CA09^+^ AM. Seven days after AM transplantation, endogenous CD45.1 SFB^-^CA09^+^ AM and different transplanted AM were FACS-sorted and challenged with live *S. pneumoniae* at MOI = 20, followed by an antibiotic protection assay. Results are displayed as CFU/mL of bacteria internalized by AM and the percentage of CFU intracellular killing. All experiments were performed with n = 3 per group. Results are shown as mean ± SD. Statistical analysis: ***p < 0.001, ****p < 0.0001, ns: not significant.

**Figure S8.**
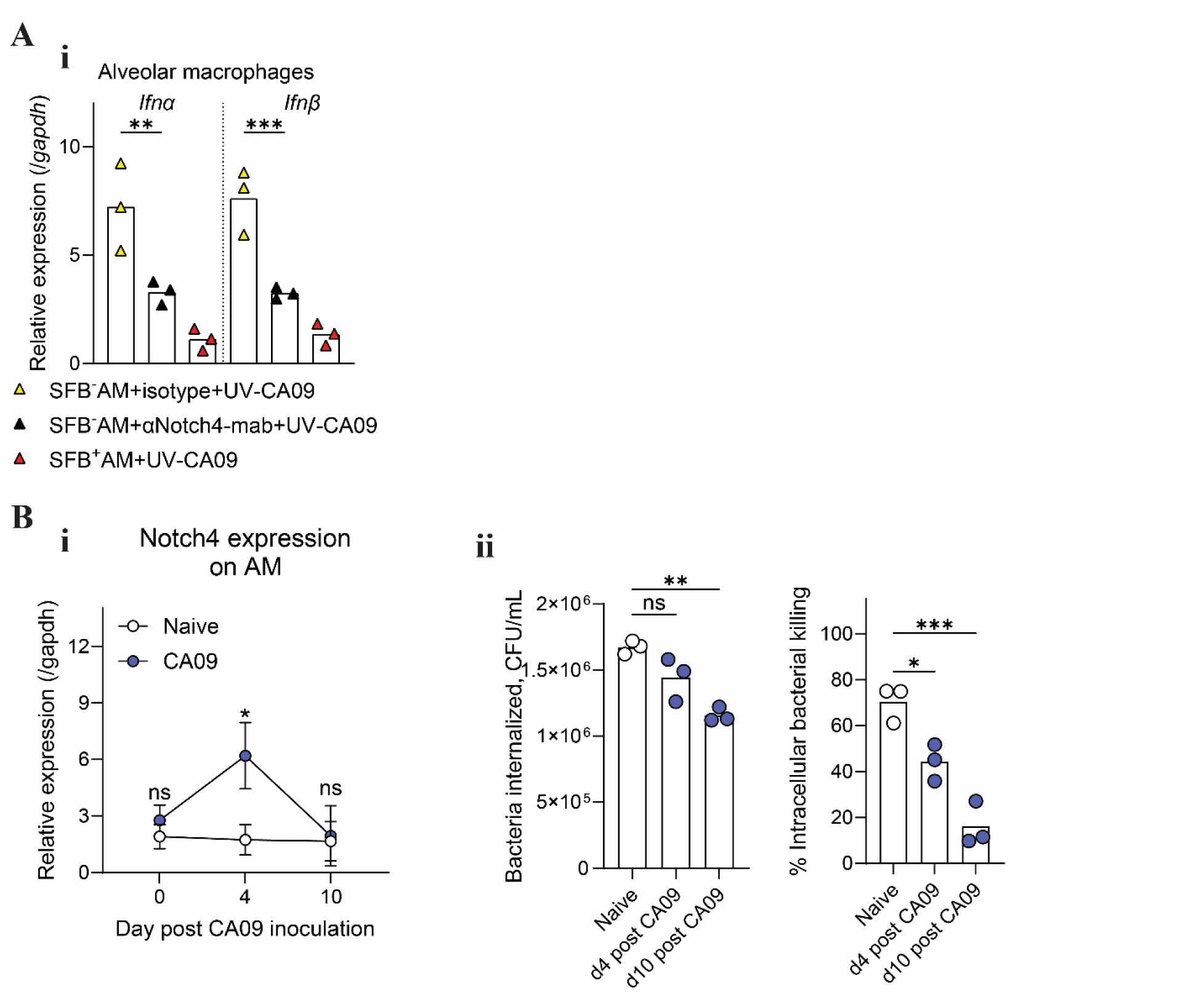
Notch4-mediated IAV-induced type I IFN associated with AM dysfunction and SFB’s alleviation. **(A)** Relative expression of *Ifn-α* and *Ifn-β* was assessed by RT-qPCR after AMs were treated with UV-inactivated CA09. **(B) (i)** Relative expression of Notch4 in AMs at days 0, 4, and 10 post-CA09 inoculation. **(ii)** bacteria internalized by AM and percentage intracellular killing by AM were quantified by an antibiotic protection assay in mice inoculated with CA09. All experiments were performed with n = 3 per group. Results are shown as mean ± SD. Statistical analysis: *p < 0.05, **p < 0.01, ***p < 0.001, ****p < 0.0001, ns: not significant.

